# Rapidly changing speciation and extinction rates can be inferred in spite of non-identifiability

**DOI:** 10.1101/2022.05.11.491456

**Authors:** Bjørn T. Kopperud, Andrew F. Magee, Sebastian Höhna

## Abstract

The birth-death model is commonly used to infer speciation and extinction rates by fitting the model to extant phylogenetic trees. Recently, it was demonstrated that speciation and extinction rates are not identifiable if the rates are allowed to vary freely over time. The group of birth-death models that are not identifiable from each other is called a congruence class. Every model in a congruence class is equally likely, and there is no statistical evidence that can be used to favor one over the other. This issue has led researchers to question if and what patterns can reliably be inferred, and whether time-variable birth-death models should be fitted at all. We explore the congruence class in the context of several empirical phylogenies as well as hypothetical scenarios and summarize shared patterns in the congruence class. We show that strong directional trends in speciation and extinction rates are ubiquitous among most models within the congruence class, and conclude that inference of strong directional trends is therefore robust. Conversely, estimates of constant rates or gentle slopes are not robust and must be treated with caution. Additionally, most conflict in trends within the congruence class is observed near the present, implying that very recent rate changes should be treated carefully. Interestingly, the space of valid speciation rates is limited in contrast to extinction rates, which are less constrained. These results provide further evidence and insights that speciation rates can be estimated more reliably than extinction rates.

## 1 Introduction

Macroevolutionary species diversification is often investigated using molecular phylogenetic trees (Ricklefs, 2007; Morlon, 2014). Phylogenetic trees are an instrumental resource for learning about diversification phenomena such as rapid radiations (Boschman and Condamine, 2022; Melo et al., 2022), key innovations (Burress and Muñoz, 2021), mass extinctions (May et al., 2016; Magee and Höhna, 2021), biogeographic drivers (Goldberg et al., 2011; Thomson et al., 2021), diversity-dependence (Etienne et al., 2012), state-dependence (Maddison et al., 2007), or environmental drivers (Condamine et al., 2019; Palazzesi et al., 2022). These applications typically use some variation of the reconstructed birth-death model (Nee et al., 1994) fitted to an extant phylogenetic tree.

Recently, Louca and Pennell (2020) demonstrated that diversification rates are not identifiable when using birth-death models fitted to extant phylogenetic trees. Given continuously differentiable rate functions, they presented a class of so-called congruent models that are equally likely. The discovery of the congruence class has led to questions of whether we should infer diversification rates at all, and if so, what inferences we can reliably make (Pagel, 2020; Louca et al., 2021; Helmstetter et al., 2022; Morlon et al., 2022). One concern is that congruence classes can contain contradictory temporal trends: for example, increasing, decreasing and flat rate functions in the same time period (Louca and Pennell, 2020). On the other hand, it is possible that some congruence classes have universal support for a specific diversification trend.

Are there diversification rate patterns, e.g., shifts or directional trends, that are shared among all models within a congruence class? Answering this question requires us to analyse the congruence class. Here we explore the congruence class using the recently developed method ACDC (Höhna et al., 2022). Specifically, we explore a variety of alternative models within the congruence class, including both specific alternative rate hypotheses and rates sampled randomly from stochastic processes. Finally, we inspect and analyse the produced alternative models for common trends.

In this study, we investigate time-varying diversification rates in a set of 286 phylogenetic clades from Condamine et al. (2019). We use the time-calibrated phylogenetic trees to estimate diversification rates under an episodic birth-death models using Bayesian inference (Magee et al., 2020), which gives one model from the congruence class. From this large group of datasets, we select three focal datasets that we believe are useful exemplars to illustrate how the congruence class behaves under certain scenarios. These scenarios include (a) constant rates, (b) a single rate-shift, and (c) a more complex history of rate increases and decreases. These three datasets, which we will discuss in detail in the main text, are: the New World/African true parrots (Psittacidae, n = 330), the tyrant flycatchers (Tyrannidae, n = 419), and the woodpeckers (Picidae, n = 223). A selection of nine additional empirical datasets can be found in the supplemental material. To capture a more complete range of possible diversification scenarios, we augmented our selection of empirical datasets with hypothetical scenarios. Specifically, we considered rates with exponential increase and decrease, sigmoidal increase and decrease (i.e., an up-shift and a down-shift), a modal burst event, and linear increase and decrease. Taken together, these empirical and hypothetical models cover a wide range of realistic diversification scenarios.

## 2 Results

### 2.1 Estimating time-varying diversification rates

We estimated time-varying diversification rates using an episodic birth-death model (Stadler, 2011; Höhna, 2015) implemented in RevBayes (Höhna et al., 2016a). We used the horseshoe Markov random field (HSMRF) distribution as prior distribution on the speciation and extinction rates to model autocorrelation among the episodes, which gives a temporal smoothing effect while allowing for larger jumps (Magee et al., 2020). This analysis provides us with a reference model with speciation and extinction rates that we will investigate further. We estimated diversification rates for 286 datasets although we only show the results of three exemplary datasets in the main text (New World/African true parrots, tyrant flycatchers and woodpeckers) and nine more examples are shown in the Supplementart Material (Figs. S1–S3).

The extinction rates for all three datasets are approximately constant (Fig. 1). The diversification history of the New World/African true parrots is relatively constant; they diversified at the same approximately constant rate throughout the entire span of the phylogeny (Fig. 1, Psittacidae). The speciation rate of the tyrant flycatchers shows a single rate shift (Fig. 1, Tyrannidae). Thus, their diversification history can be split in three parts: first a period of constant low diversification until around 45 Ma, second a short period of a sharp increase in speciation rate, and third another relatively constant but high period of diversification starting at 40 Ma and continuing towards the present (Fig. 1, Tyrannidae). The inferred speciation rate of woodpeckers depicts a more complex diversification rate scenario with increases and decreases. The speciation rate inferred using the woodpeckers dataset exhibit an initial speciation rate increase around the time the wrynecks (Jynginae) and the piculets (Picumninae) split off (25 Ma, Fig. 1, Picidae). After that, there is a constant period (25-18 Ma), followed by another increase in speciation rate corresponding to the radiation of the true woodpeckers (Picinae). After peaking at around 8 Ma, the woodpecker speciation rate decreases towards the present. The three exemplary datasets show three qualitatively distinct patterns: constant vs. single shift vs. complex diversification.

**Figure 1:**
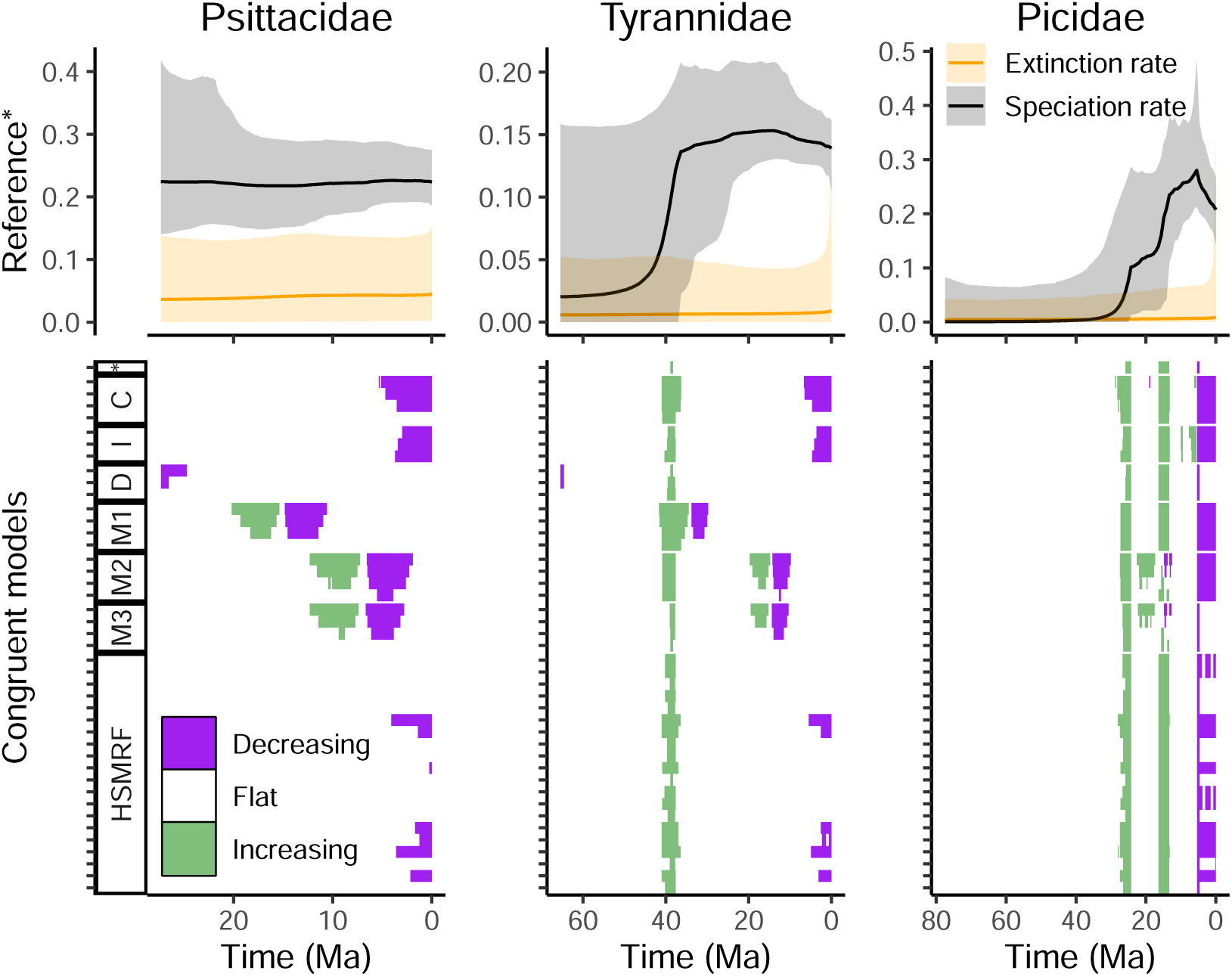
A summary of the congruence class for a set of different datasets: New World/African true parrots (Psittacidae), tyrant flycatchers (Tyrannidae), and woodpeckers (Picidae). The top row depicts the posterior median and 95% credible interval speciation and extinction rates, for the episodic birth-death model implemented in RevBayes. The bottom row depicts summaries of directional trends in the congruence class, where each row is a model. The models include alternative extinction rates that are constant (C), exponentially increasing (I), exponentially decreasing (D), modal (M1-M3), HSMRF-distributed (horseshoe Markov random field), and the reference model (*). If the slope of the speciation rate function is greater than or less than *ϵ* = 0.02 rate units per million years, we say that the function is increasing, or decreasing, respectively. An additional nine datasets can be found in Figs. S1–S3, and the exact rate shapes of the alternative models are in Figs. S4–S6.

### 2.2 Alternative speciation rates may be strongly constrained

We used the R-package ACDC (Höhna et al., 2022) to explore the congruence class, i.e., to compute the pulled diversification rates and to propose alternative congruent models. ACDC takes in a proposed alternative speciation (or extinction) rate, and uses the pulled diversification rate to determine the extinction (or speciation) rate necessary to remain in the congruence class.

We made a detailed examination of alternative models for the New World/African true parrots, tyrant flycatchers and woodpeckers (Fig. 1). Using a number of alternative rate functions, we proposed both new extinction rates and new speciation rates. We found that proposing alternative speciation rates is more difficult than proposing alternative extinction rates. When we proposed alternative speciation rates for the 12 empirical data sets, we inferred extinction rate trajectories that included negative rates for 76% of the models. For the hypothetical scenarios, 36% of the proposed speciation rates resulted in negative extinction rate functions. Negative extinction rates do not make sense mathematically or biologically. This implies that many of the proposed speciation rate functions are not valid choices. Conversely, when we proposed alternative extinction rates, we never obtain invalid (i.e., negative) alternative speciation rates function neither for the empirical nor for the hypothetical scenarios. Thus, it appears that any extinction rate function is contained within the congruence class but not all speciation rate function. In the next sections, we focus on proposing new extinction rates rather than new speciation rates.

### 2.3 Rapid changes are shared among congruent models

We wish to compare shared features across the models, which raises the question, how do we compare features among birth-death models? Biologists are often interested in whether the rates increased, decreased, or remained constant. Thus, we investigated if the directional trends were significantly increasing or decreasing within a short time period. First, we show the significant directional trends for the New World/African true parrots (Psittacidae), the tyrant flycatchers (Tyrannidae), and the woodpeckers (Picidae). Then we show the significant directional trends for the hypothetical scenarios.

We explored the congruence class for three empirical datasets (Fig. 1). We used a variety of alternative extinction rate trajectories to represent a wide range of possible extinction scenarios (Figs. S4–S6). The significant directional trends in the speciation rates are consistent across the congruence class when the slope of the speciation rate is steep (Fig. 1, Tyrannidae and Picidae). Specifically, we were able to recover the following events. The single rate shift in the tyrant flycatchers is unanimously represented in our sample of the congruence class (Fig. 1, middle column, bottom plot). For the woodpeckers, we were able to recover two events where the speciation rate increased, and another event where the speciation rate decreased (Fig. 1, right column, bottom plot). When the speciation rate is relatively constant or shallow, then there is no consistent directional trend in the speciation rate across the congruence class. In other words, when the rates are constant, it is trivial to propose an alternative congruent model that contradicts the diversification pattern of the reference model. These results are corroborated by an additional nine empirical datasets that we present in the Supplementary material (Figs. S1–S3).

Despite our initial analyses covering 286 clades, there were only a limited set of interesting scenarios. For example, we never inferred speciation rates that were monotonously decreasing across the time span of the phylogeny, or that resembled exponential growth or decline. To complement our analysis of the behaviour of the congruence class using empirical datasets, we added hypothetical scenarios with more variable shapes. These scenarios are commonly used in diversification rate analyses (see Morlon et al. 2016, Höhna 2014, Höhna et al. 2016b, and Magee et al. 2020). Specifically, we assessed a shallow linear increase, an exponential decrease, and a sigmoidal decrease (Fig. 2). The assessments from the hypothetical scenarios are similar to the empirical scenarios. A shallow change in the speciation rate is not well recovered by the inferred rates in the congruence class (L+ and E–, Fig. 2). The steep descent of the sigmoidally decreasing model is recovered by our sample of the congruence class. Unlike the empirical datasets, however, this descent is not precisely overlapping in time. The proposed models exhibit a decrease in speciation rate between 30 and 50 Ma, but with some temporal distortion (Fig. 2, S–). In the supplemental material, we also investigated hypothetical models where the extinction rate has a distinct pattern, and we explored the congruence class by proposing alternative speciation rates. Similar to speciation rates, we show that rapidly changing extinction rates are also robust to the congruence class (Fig. S25–S32).

**Figure 2:**
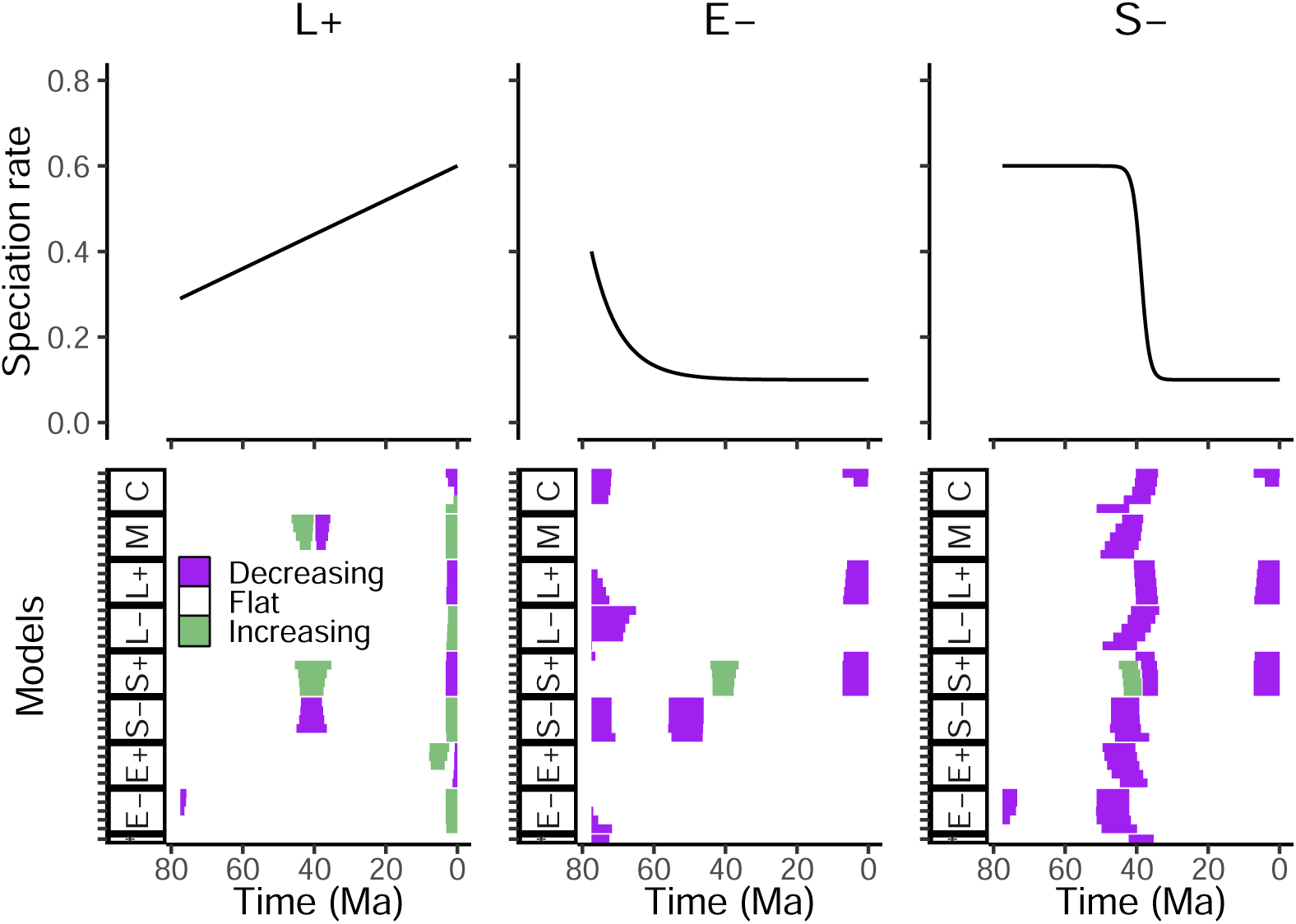
Three hypothetical scenarios: a linear increase in speciation rate, an exponential decrease, and a sigmoidal decrease. The reference model is the speciation rate in black, and a constant extinction rate of *µ* = 0.28. For each congruence class, we proposed various alternative extinction rates: constants (C), modals (M), sigmoidal increase (S+), sigmoidal decrease (S-), linear increase (L+), linear decrease (L-), exponential increase (E+) and exponential decrease (E-), with five rate trajectories for each of these shape categories (Fig. S15). Each congruence class results in a set of models with inferred speciation rate curves (Figs. S19, S20, S23). We used a threshold of *ϵ* = 0.02 rate units per Ma to compute the significant directional trends in the inferred speciation rates. We selected a time scale similar to the woodpecker phylogeny for interpretability.

For both the empirical and hypothetical scenarios, we observed the strongest disagreement of trends in the intervals near the present (Figs. 1 and 2, and Fig. S15). Speciation rate changes near the present are strongly driven by the constraint that all speciation rate functions must be equal at the present. By choosing higher alternative extinction rates than the reference extinction rate, we obtain higher alternative speciation rates for most of the evolutionary history which then must drastically drop towards the present to *λ*_0_ (Figs. S4–S6). Conversely, lower alternative extinction rates than the reference extinction rate leads to lower speciation rates until the present where the speciation rate rapidly increases towards *λ*_0_. Thus, abrupt rate change in the speciation rate near the present may entirely be an artefact of the congruence class.

### 2.4 The effect of diversification-rate uncertainty is substantial

Thus far we have explored the congruence class and investigated the robustness of diversification rate patterns of a single point estimate of the rates for each episode, either the posterior median or a hypothetical rate. However, we must acknowledge there is substantial uncertainty in the estimation of the speciation and extinction rates (Fig. 1, top row; Fig. S10). This uncertainty in the timing of the rate shifts motivated us to test different window sizes to assess whether rates have changed within a larger time span (Fig. S11).

Recall that the posterior median trend summaries had some unambiguous trends: the three rapid changes in the woodpeckers dataset and the single speciation rate increase in the flycatcher dataset (Fig. 1). When we instead study the congruence class and estimation uncertainty together, then the signal of the three intervals is only partially present (Fig. 3). Thus, the diversification rate trends are no longer unambiguous. The assessment of the rate shift is sensitive to the window size, with too small and too large windows not recovering the trend (Fig. S11). It is apparent from the posterior samples that no patterns are fully covered, and none are supported even by half coverage. Our results indicate that the uncertainty in extinction and speciation rate estimation is by far a greater problem for diversification inference than the issue of non-identifiable rates.

**Figure 3:**
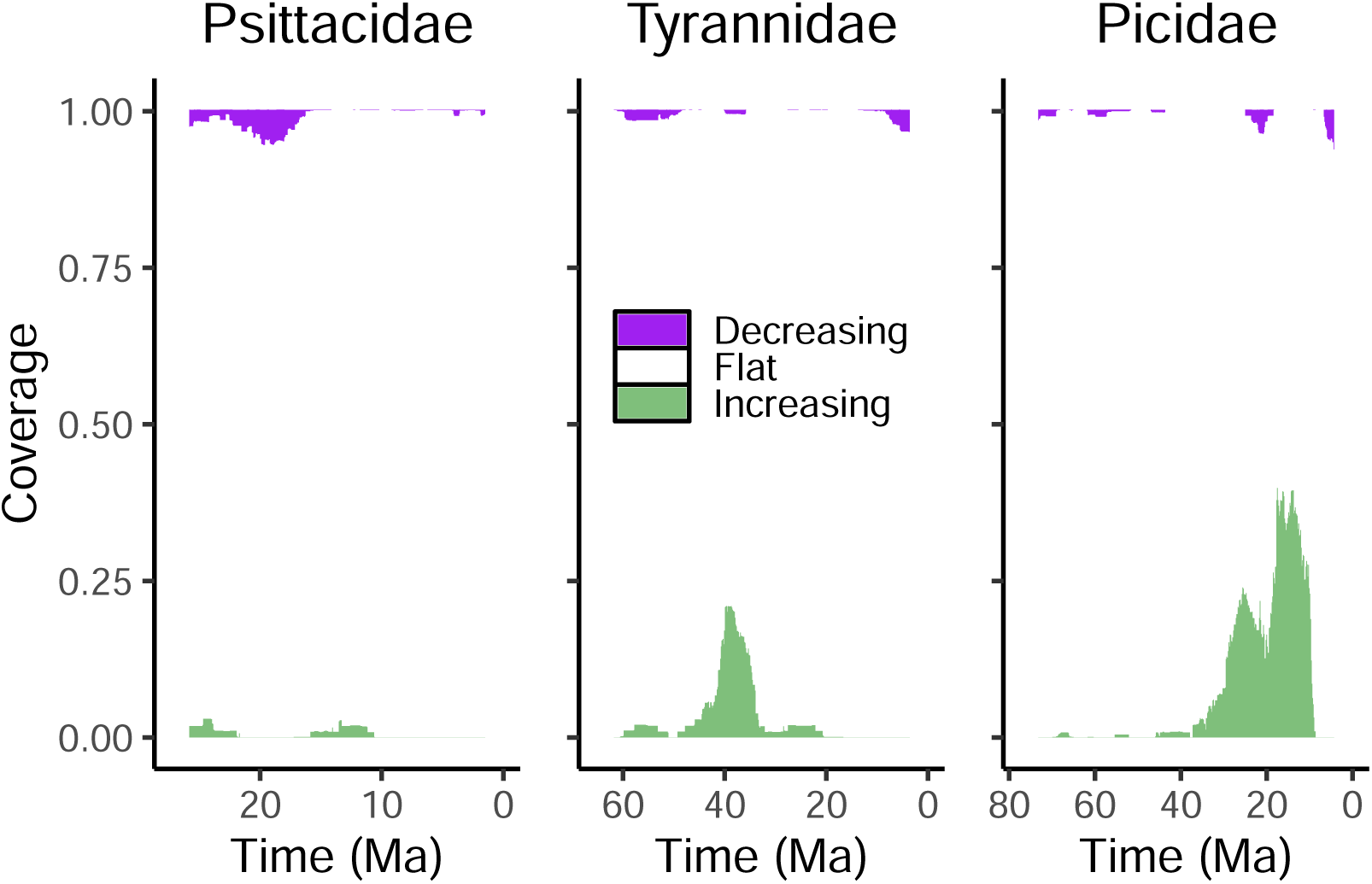
The interplay between the variation of rates in the posterior distribution and the congruence class. Each panel represents 110 samples from the posterior distribution. These samples are not congruent. For each of the samples, we constructed an additional 10 congruent models with horseshoe-distributed extinction rates. The y-axis is sorted, meaning each row does not correspond to a single model. We calculated the slopes using a window size of 55 (Δ*λ*_*i*_ = (*λ*_*i*_ − *λ*_*i*−55_)/(*t*_*i*_ − *t*_*i*−55_)). For other window sizes see Fig. S11.

## 3 Discussion

### 3.1 Conflict in rates near the present

We observed the strongest conflicts in trends near the present. The value of the speciation rate at the present is fixed to *λ*_0_ for all models within the congruence class. This means that if we are able to infer the correct congruence class, then we are also able to infer the correct speciation rate at the present. However, the trend of the speciation rate towards the present is entirely determined by the choice of the extinction rate in the near present. A greater assumed extinction rate near the present results in greater a speciation rate near the present and thus a decline towards the present; while a smaller assumed extinction rate near the present results into a smaller speciation rate near the present and thus an increase towards the present. Therefore, we should not trust the trends of the inferred diversification rates in the near present.

The common expectation is that diversification rates can be estimated more robustly towards the present compared with the more ancient half of the evolutionary history of the study group (see, e.g., May et al., 2016). There is more information due to more speciation events close to the present, which is often seen in smaller uncertainty (i.e., credible intervals) for diversification rates close to the present. Unfortunately, this higher precision might be completely artificial. Instead, we are only better able to estimate the congruence class at the present but the congruence class itself shows contradictory trends in the near present.

This observation has also strong implications for methods relying on tip rates (Freckleton et al., 2008; Jetz et al., 2012; Quintero and Jetz, 2018; Title and Rabosky, 2019). If our ability to estimate the correct trend is worse near the present, then tip rates, which are by definition near the present, are most impacted by the congruence class.

### 3.2 The space of valid speciation rates versus extinction rates

We observed that any proposed alternative extinction rate resulted in a valid, i.e., non-negative, speciation rate function. Conversely, alternative speciation rate functions must be constructed carefully to avoid nonsensical results such as negative extinction rate values. First, the space of possible speciation rate functions is by definition more restricted because the speciation rate at present, *λ*(*t*=0) = *λ*_0_, is the same for all models, whereas there is no such restriction for extinction rates. This implies that there is only one possible model with a constant speciation rate, but infinitely many possible models with constant extinction rates. Second, for any chosen alternative extinction rate we can compute a non-negative speciation rate. However, the reverse is not true. When we chose an alternative speciation rate that was smaller than the reference rate, this sometimes led to negative extinction rates (see Figs. S26–S32). Third, mismatching slopes between the reference speciation rate and the alternative speciation rate result in amplified extinction rate changes (see Figs. S25–S32). This phenomenon has also been observed by Louca and Pennell (2021) where the alternative model within the congruent class had a negative extinction rate.

On the other hand, even if not every possible alternative speciation rate function is valid, there are still infinitely many valid speciation rate functions. That is, while we can not specify any arbitrary alternative speciation rate function and obtain a sensible alternative extinction rate function, we can specify any arbitrary alternative extinction rate function and obtain a valid alternative speciation rate function. Thus, there are infinitely many speciation rate functions within the congruence class but they are very likely clustered in a small region of the space of possible rate functions.

We conclude that the space of valid alternative speciation rates is much narrower than the space of valid alternative extinction rates. This supports previous evidence that speciation rates are more reliably estimated than extinction rates from molecular phylogenies (e.g., Maddison et al., 2007; Pyron and Burbrink, 2013; Höhna, 2014; Title and Rabosky, 2019). Furthermore, since in all our examples the pure-birth model was contained in the congruence class, inferring the congruence class using a pure-birth model may be a better approach (see also Helmstetter et al. 2022).

### 3.3 Posterior distribution

We have assessed the congruence class issue primarily by using a single diversification rate (the posterior median, since we used Bayesian inference). Other lines of research that use maximum likelihood to fit the model also typically result in a single rate trajectory. Both the posterior median and the maximum likelihood estimate are point predictions of the model fit; neither indicate whether the rate estimates are precise or uncertain. We have demonstrated that the uncertainty induced by credible intervals is substantial in our selection of models. The rate uncertainty can also be substantial for models inferred using maximum likelihood, even if the uncertainty is not explicitly investigated as is often the case.

Our results indicate that the variation of rates induced by the congruence class is, in a sense, much less than the variation represented by the inherent uncertainty in the time-varying model. In other words, we argue the the congruence class is a smaller problem compared to the uncertainty in rate estimates of time-varying models. The inferred trends were robust to the congruence class but less so to the posterior uncertainty. If a shift was present in the reference model, then all models in the congruence class agreed on the same shift unambiguously, whereas the posterior samples varied in the timing of the rate shifts.

One of the main challenges for inferring trends when including estimation uncertainty is the timing of the rapid rate shift. Often, there is large uncertainty of the exact timing of the rate shift but less uncertainty whether the shift occurred. However, our current trends analysis is focusing on whether a trend occurred at a given time interval, and therefore did not agree among all diversification rate samples. This problem is potentially more severe when phylogenetic and divergence time uncertainty is included in the assessment of diversification rate patterns. Instead of our test for a rate change at a given time interval one might instead ask if there was at least one rate change (or how many such rate shifts there were).

## 4 Concluding remarks

Non-identifiability of diversification rates is a real issue and should be addressed when studying temporal patterns in diversification histories. Yet, the congruence class may not be a problem when testing for macroevolutionary diversification hypotheses. Based on our selection of datasets and alternative models, we argue that abrupt changes in speciation and extinction rates are robust to the congruence class. Conversely, periods of constant or flat speciation and extinction rates are not robust, and it is trivial to construct congruent but contradictory evolutionary hypotheses. Furthermore, the restriction of the congruence class that all models share the same speciation rate at the present *λ*_0_, often induces diversification rate changes near the present. These rate shifts near the present are likely to be artefacts and should be treated cautiously. Additionally, we conclude that the space of possible speciation rate functions is much more restricted than the space of possible extinction rate functions. Indeed it appears from our albeit limited exploration that essentially any extinction rate function is contained within the congruence class, i.e., we were never able to propose an extinction rate function that was invalid. However, when we proposed alternative speciation rate functions we often obtained invalid extinction rate functions, which implies that not all speciation rate functions are valid. Therefore, the space of possible speciation rate functions is narrower within the congruence class which provides reason to be optimistic that patterns of speciation rates are robust despite the congruence class.

## 5 Methods

### 5.1 Empirical datasets used as case studies

The empirical phylogenetic trees were obtained from Condamine et al. (2019). Specifically, the three highlighted bird families were derived from the study by Jetz et al. (2012). In our study, we used a single phylogeny as the observation, thus treating the phylogenetic trees as known without error. We also knew the proportion of taxa sampled (*ρ*) as the fraction of tips divided by the number of described taxa. This sampling fraction was directly used in our study to correct for uniform incomplete taxon sampling (Höhna et al., 2011).

### 5.2 Estimation of the reference diversification model

We fitted a time-varying birth-death process to the empirical trees in RevBayes (Höhna et al., 2016a). Evidence suggests that it is possible to estimate the correct congruence class, i.e., any model that is congruent to the true model and thus has the same likelihood as the true model (Magee et al., 2020), if not the true model when using episodic models (Legried and Terhorst, 2021). We treated the rate functions as piecewise constant (100 pieces), and we used a horseshoe Markov random field (HSMRF) process to specify a prior distribution on the speciation and extinction rates. We followed Magee et al. (2020) when specifying the parameters of the HSMRF distribution.

Specifically, we assumed the following prior distribution for the extinction rate *µ*_0_ at the present and all consecutive extinction rates *µ*_*i*_:

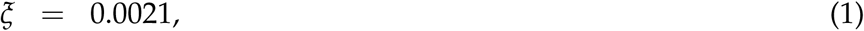

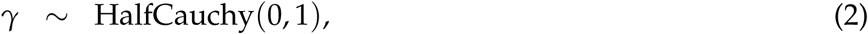

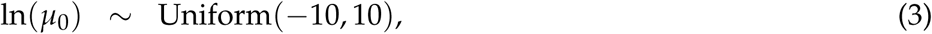

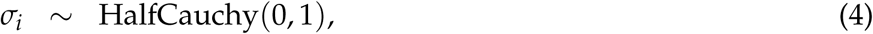

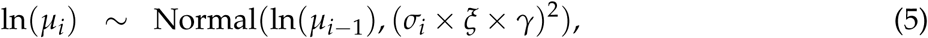

where *ξ* scales the overall magnitude of the changes in accordance with the number of episodes (here 100), and *σ*_*i*_ is the local scaling factor. We picked *ξ* = 0.0021 such that we expected *a priori* ln(2) shift events of magnitude two or higher. An exactly matching set of prior distributions were assumed for the speciation rates.

We used `RevBayes` to fit the model in a Bayesian framework, and we used the Metropolis-Hastings algorithm to sample from the posterior distribution. We ran four chains in parallel for 50,000 iterations, discarded the first 5,000 iterations as burnin, and compared the chains using the Kolmogorov-Smirnov test to test for convergence (Fabreti and Höhna (2022), see Supplementary materials). We used the posterior median estimates for extinction rate and speciation rate for further analyses.

### 5.3 Exploring the congruence class

We imported the fitted time-varying birth-death models into the R-package `ACDC`: Analysis of Congruent Diversification Classes (Höhna et al., 2022). We used the pulled net-diversification rate, *r*_*p*_, to construct the congruence class (Louca and Pennell, 2020):

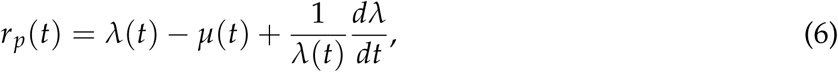

where *t* is in units of time before present. `ACDC` uses numerical approximations, i.e., piecewise linear rate functions, to solve the derivative and to construct the congruence class. We used a grid of 500 time points for the piecewise approximations, which has been shown to give precise approximations of the underlying continuous functions (Höhna et al., 2022).

In order to construct an alternative model in the congruence class, `ACDC` requires an alternative extinction or speciation rate function. Then, `ACDC` solves Eq. (6) for either *λ* or *µ*, depending on which alternative rate function is proposed, to compute the corresponding alternative extinction or alternative speciation rate function, respectively. We detail how the alternative rate functions are proposed in the next sections.

#### 5.3.1 Specific diversification hypotheses

We employed two strategies to sample alternative models from the congruence class, by generating alternative extinction or speciation rates. First, we selected a set of specific hypotheses that are biologically distinct. For the extinction rates, we constructed constant rate functions, increasing rate functions, decreasing rate functions, and a scenario in which there is a short-lived increased (modal) in extinction rate (Figs. S4–S6). Similarly, we constructed a set of alternative speciation rate functions, including linear increasing speciation rates, and an abrupt up-shift or down-shift (Figs. S4–S6).

#### 5.3.2 Drawing random diversification rates

Our second approach to exploring the congruence class was to sample rate functions from a random probability distribution. Specifically, we used the horseshoe Markov random field (HSMRF) (Carvalho et al., 2010; Magee et al., 2020) distribution to model temporally autocorrelated rates. First, the simulation starts by drawing the global variance parameter *γ* ∼HalfCauchy(0, 1). Then, for the case of simulating extinction rates, we draw *µ*_0_ ∼lognormal 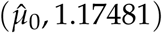, where the index 0 is at the present and 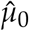 is the estimated extinction at the present for the reference model. In the case of simulating speciation rates, we explicitly set 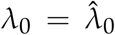 because the congruence class requires that all alternative models have the same speciation rate at the present.

For each preceding episode *i* we draw both a new local scaling factor *σ*_*i*_ ∼HalfCauchy(0, 1) and then a new rate ln(*µ*_*i*_) ∼Normal(ln(*µ*_*i*_ −_1_), (*σ*_*i*_ ×*ξ*× *γ*)^2^), where *ξ* scales the overall magnitude of the changes in accordance with the number of episodes (here 100), such that we expected *a priori* ln(2) shift events of magnitude two or higher. This results in a distribution of ln(*µ*_*i*_) that is centered around the previous ln(*µ*_*i*−1_), but with fat tails on each end. The most probable draw is little to no change, but large jumps are also possible.

### 5.4 Sampling congruent models from the posterior

We sampled 110 rate functions from the posterior distribution to represent the uncertainty in the parameter estimates. The samples were equidistant to minimize MCMC autocorrelation. Each of these diversification rate samples have different likelihoods, meaning they are not congruent. The individual posterior samples have more rapidly and sporadically changing rates, as opposed to the relatively smooth median summaries (Fig. S10).

For each of these 110 samples, we constructed the congruence class and sampled 10 alternative models using HSMRF-distributed extinction rate functions. We fixed the initial episode (*µ*_0_) of the proposed extinction rates to being equal to the posterior sample initial extinction rate. This avoids extra artificial bias in directional trends near the present, since the speciation rate must always be equal at the present (*λ*_0_). If we had chosen the same value for *µ*_0_ for all posterior samples, then the directional trends in speciation rate in the near present (Δ*λ*) would be completely determined by the first episode in the posterior samples.

For large proposed extinction rates, we used rejection sampling to ensure that the maximum of the proposed rate was not greater than the maximum of *µ*_0_ + 0.1 across the posterior samples (woodpeckers: 0.32, flycatchers: 0.62, parrots: 0.28). For some posterior-congruent models, we encountered numerical issues preventing us from computing the corresponding speciation rate, and we consequently discarded these samples.

We assessed the common trends in the speciation rate functions in the resulting models (Fig. 3). Since the number of models are now large, we sorted the model axis such that the “increasing” and “decreasing” bars are stacked. This means that one row no longer corresponds to a single model. However, we can still assess the fraction of significant trend directions for a particular time interval.

### 5.5 Assessing significant rate changes

We assessed common patterns in diversification rates by computing the trends in diversification rate changes. Specifically, we computed the slope of the speciation rate function, as Δ*λ*_*i*_ = (*λ*_*i*_ − *λ*_*i* − *k*_)/(*t*_*i*_ − *t*_*i* − *k*_), where the window size *k* is unity or some larger value (see Fig. S11). If the slope is greater than some threshold *e*, we interpret the rate function as increasing. If the slope is smaller than −*ϵ*, we interpret the trend as decreasing. If the difference is between *ϵ* and −*ϵ*, we interpret it as flat.

The selection of *ϵ* is to some extent arbitrary, and therefore we explored three values for *ϵ* ∈ {0.01, 0.02, 0.05}. Choosing a too large *ϵ* will result in no significant changes being detected at all (Fig. S9). Conversely, too small *ϵ* will result in small noisy changes being detected as significant changes (Fig. S7). In our analyses, we mainly used *ϵ* = 0.02 rate units per Ma as the threshold.

### 5.6 Specifying and exploring hypothetical scenarios

We constructed a set of specific scenarios that we deemed either biologically realistic or interesting to complete the empirical exploration of the congruence class. Specifically, we assessed constant rates, a modal burst event, linearly increasing and decreasing, an up-shift and a down-shift, as well as exponentially increasing and decreasing rates. For each scenario, we selected one reference speciation rate, and used in all cases a constant reference extinction rate of *µ* = 0.28. Further, we used all of the proposed rate functions as proposed alternative extinction rate functions to explore the congruence class. Each individual rate function can be found in the Supplementary materials, in Figs. S16-S23.

## 6 Data and code availability

The data and code necessary for reproducing the analyses and the figures is available on github (http://github.com/kopperud/cc_exploration).

## 8 Author contributions

BTK and SH conceived the study. All authors contributed to methods development. BTK performed the analyses. BTK and SH drafted the manuscript. All authors revised and approved the manuscript.

