## Supplemental Information for "Rapidly changing speciation and extinction rates can be inferred in spite of non-identifiability"

#### Supporting information

Bjørn T. Kopperud[1,2], Andrew F. Magee[3] and Sebastian Höhna[1,2*]

[1] *GeoBio-Center, Ludwig-Maximilians-Universität München, Richard-Wagner Straße 10, 80333 Munich, Germany*
[2] *Department of Earth and Environmental Sciences, Paleontology & Geobiology, Ludwig-Maximilians-Universität München, Richard-Wagner Straße 10, 80333 Munich, Germany*
[3] *Department of Biostatistics, University of California, Los Angeles, 90095, U.S.A.*

### S1 Extended datasets

In this section we plot the estimated reference model and summary of the directional trends for an additional nine datasets. These datasets provide more empirical examples about the robustness of estimated diversification patterns and trends within the congruence class. Overall, the results corroborate our main findings in the main text: (1) sharp diversification rate changes are robust and unambiguously supported, (2) trends in the near-present show the strongest conflicting signal, and (3) reference models with comparable flat diversification rates provide ambiguous signal within the congruence class.

We chose to first include the finches (Fringillidae), manakins (Pipridae), and the most species-rich family of tubenose birds (Procellariidae) (Fig. S1). In Fig. S2 we depict the equivalent congruent models for another three families: mice (Muridae), white-eyes birds (Zosteropidae), and the evening primroses (Onagraceae). The diversification histories of the indigobirds, whydahs, and the cuckoo-finch (Viduidae), the New World warblers (Parulidae), and free-tailed bats (Molossidae) are comparatively more recent and their rates change more abruptly (Fig. S3). For this reason, we assessed the directional trends with a more conservative threshold of $\epsilon = 0.08$ rate units per million years (Fig. S3). The tree for the evening primroses was taken from Freyman and Höhna 2019, while the remaining trees were taken from Condamine et al. 2019.

The significant directional trends in the speciation rates are consistent across the congruence class when the slope of the speciation rate is steep (Figs. S1 to S3). When the speciation rate is shallow, it is trivial to construct a model that is in disagreement with the reference model.

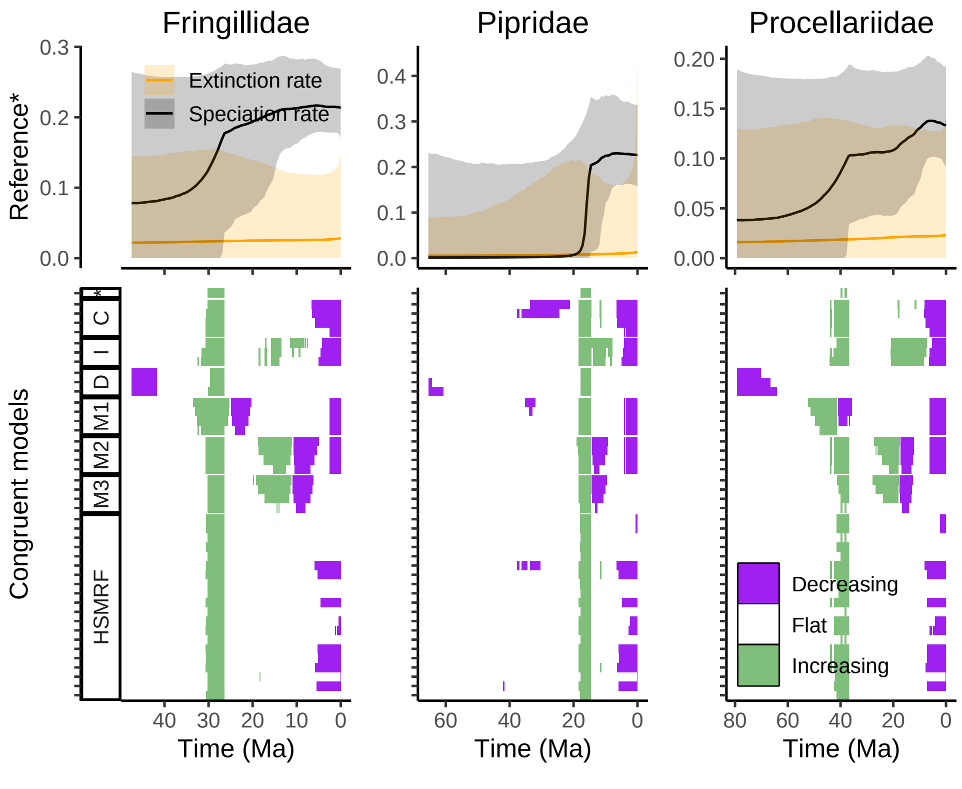

**Figure S1:** A summary of the congruence class for three bird families: finches (Fringillidae), manakins (Pipridae), and the most species-rich family of tubenoses (Procellariidae). The top row depicts the posterior median and 95% credible interval speciation and extinction rates, for the episodic birth-death models implemented in `RevBayes`. The bottom row depicts summaries of directional trends in the congruence class, where each row is a model. The models include alternative extinction rates that are constant (C), exponentially increasing (I), exponentially decreasing (D), modal (M1-M3), HSMRF-distributed (horseshoe Markov random field), and the reference model (*). If the slope of the speciation rate function is greater than the threshold $\epsilon$ (in rate units per million years), or less than $-\epsilon$, we say that the function is increasing, or decreasing, respectively. We used a threshold of $\epsilon = 0.01$ for the finches and manakins, and $\epsilon = 0.005$ for the tubenoses.

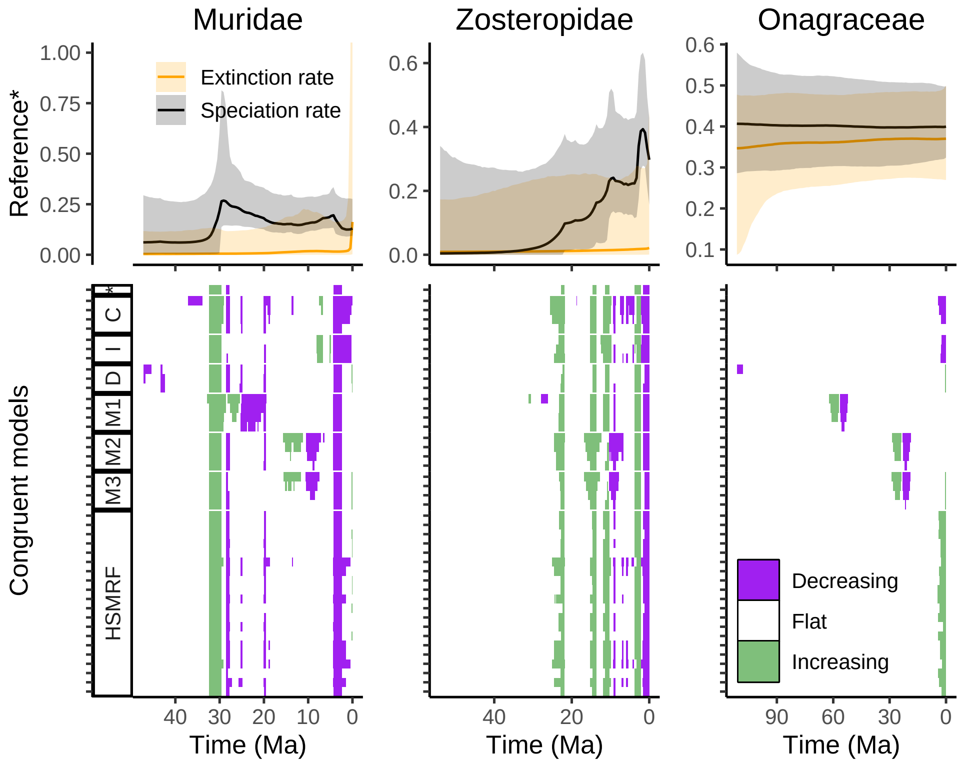

**Figure S2:** A summary of the congruence class for three families: mice (Muridae), white-eyes birds (Zosteropidae), and the evening primroses (Onagraceae). The top row depicts the posterior median and 95% credible interval speciation and extinction rates, for the episodic birth-death models implemented in `RevBayes`. The bottom row depicts summaries of directional trends in the congruence class, where each row is a model. The models include alternative extinction rates that are constant (C), exponentially increasing (I), exponentially decreasing (D), modal (M1-M3), HSMRF-distributed (horseshoe Markov random field), and the reference model (*). If the slope of the speciation rate function is greater than $\epsilon = 0.02$ (in rate units per million years), or less than $-\epsilon$, we say that the function is increasing, or decreasing, respectively. For the evening primroses, we added a constant of 0.3 such that $\mu'(t) = \mu(t) + 0.3$ for all proposed extinction rates. This accommodates their overall higher diversification rates, and prevents artifacts in the near-present due to the congruence class requiring $\lambda_0$ to be equal across all models.

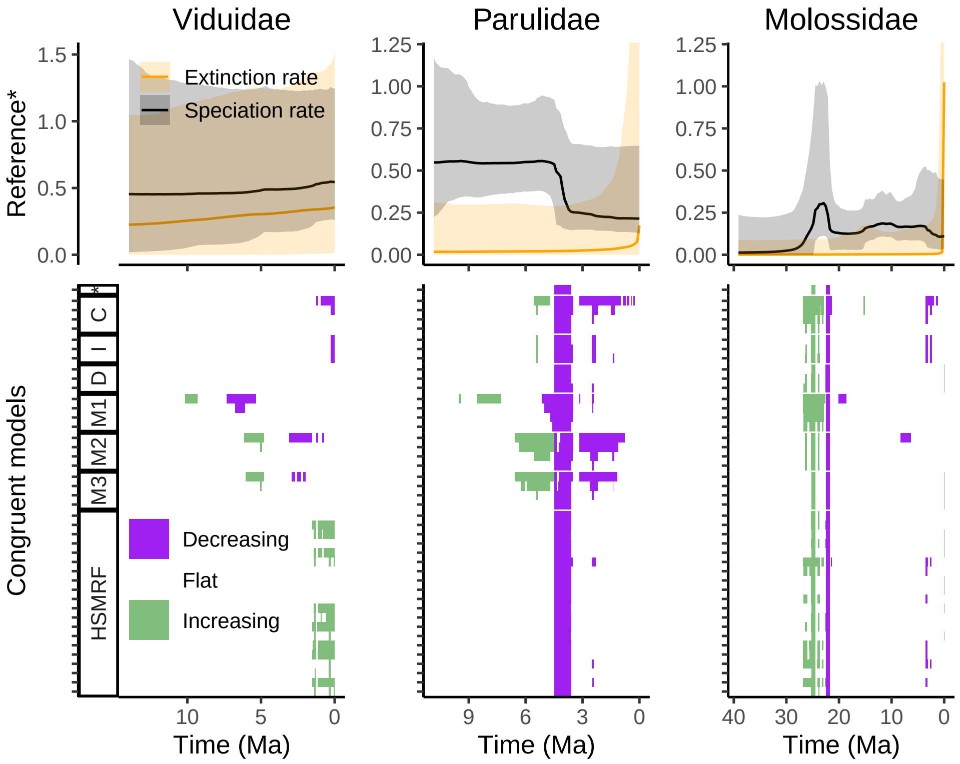

**Figure S3:** A summary of the congruence class for three families: indigobirds, whydahs, and the cuckoo-finch (Viduidae), the New World warblers (Parulidae), and free-tailed bats (Molossidae). The top row depicts the posterior median and 95% credible interval speciation and extinction rates, for the episodic birth-death models implemented in `RevBayes`. The bottom row depicts summaries of directional trends in the congruence class, where each row is a model. The models include alternative extinction rates that are constant (C), exponentially increasing (I), exponentially decreasing (D), modal (M1-M3), HSMRF-distributed (horseshoe Markov random field), and the reference model (*). If the slope of the speciation rate function is greater than $\epsilon = 0.08$ (in rate units per million years), or less than $-\epsilon$, we say that the function is increasing, or decreasing, respectively. For the Viduidae, we added a constant of 0.25 such that $\mu'(t) = \mu(t) + 0.25$ for all proposed extinction rates. This accommodates their overall higher diversification rates, and prevents artifacts in the near-present due to the congruence class requiring $\lambda_0$ to be equal across all models.

#### S2 Detailed exploration of the congruent models for the three main empirical datasets

In this section we plot the extended constructed congruent models set for the for the New World/African true parrots (Fig. S4), the tyrant flycatchers (Fig. S5), and the woodpeckers (Fig. S6). That is, we show the alternative rate functions within the congruence to provide a more detailed picture of the congruence class. These alternative rate functions were used in the main text to assess the robustness of the inferred diversification rate trends.

Most of the woodpecker alternative models exhibit the same three major trends in the woodpecker speciation history: an increase around 25 Ma, and another increase around 18 Ma, and a decrease starting around 8 Ma (Fig. S6). All alternative models within the congruence class agree on the overall signal of two increases, which is consistent with previous findings of a burst in speciation for the true woodpeckers (Picinae), and a later increase for the piculets (Picumninae) (Shakya et al., 2017). Most alternative models show no other significant trends, with the exception of the modal models. For those, we were able to induce a short-lived increase, followed by a short-lived decrease in speciation rate (Fig. S6, M2-M3). Similarly, in the diversification rates of the flycatcher dataset, there is one strong rate shift at around 40 Ma, which is unanimously recovered by the congruence class (Fig. S5). The parrot diversification history exhibits relatively constant rates and the alternative models showed conflicting trends (Fig. S4). When the rates are constant, it is trivial to propose an alternative congruent model that contradicts the diversification pattern of the reference model.

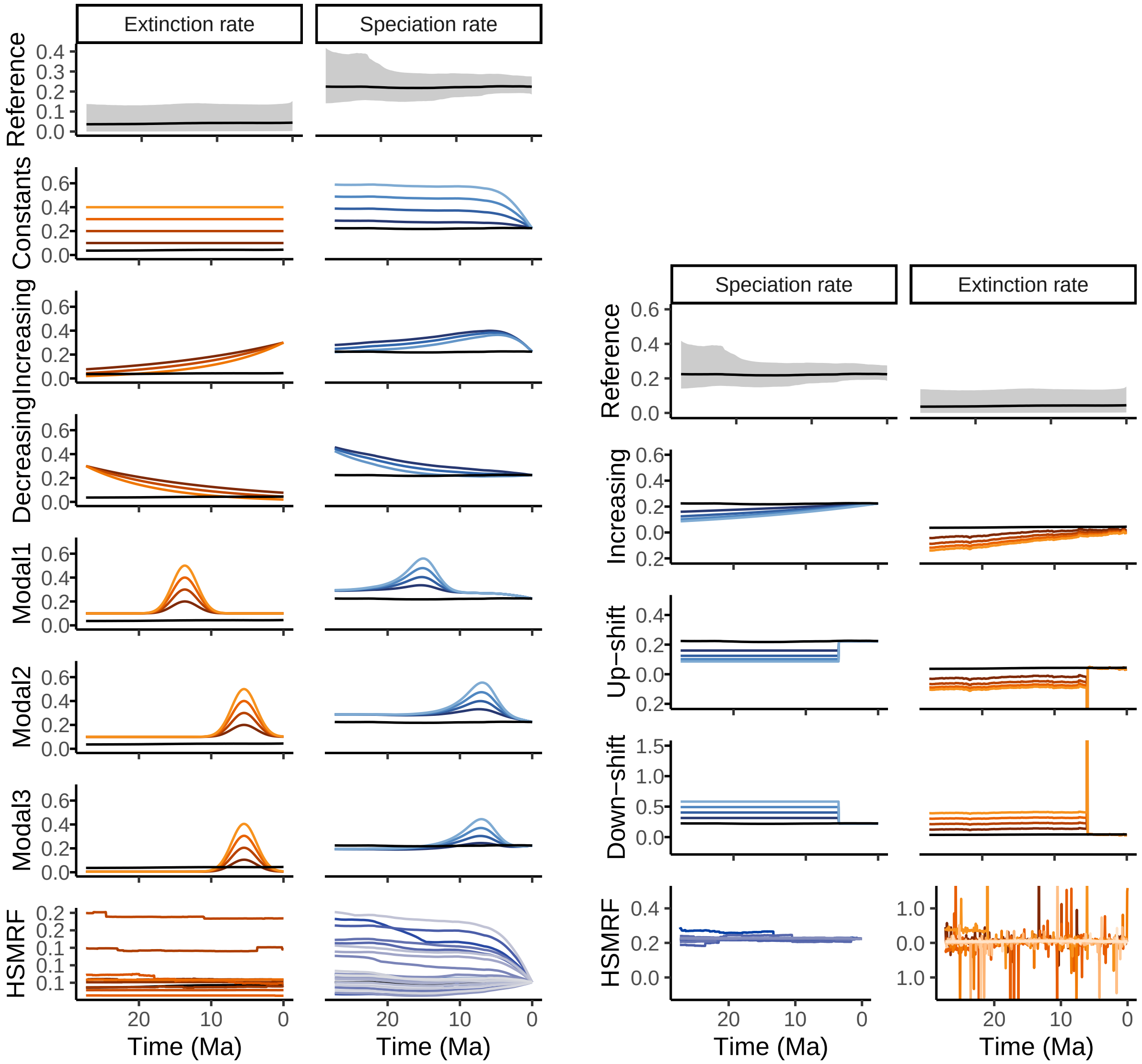

**Figure S4:** Congruent birth-death models for the New World/African true parrots (Psittacidae), based on a time-calibrated phylogenetic tree from (Condamine et al., 2019). Top row: the episodic birth-death model fitted in `RevBayes`, where the posterior median is in black, and the 95% credible interval in grey. The left panel exhibits models that are constructed by proposing alternative extinction rates. The right panel exhibits models that are constructed using alternative speciation rates. Each color-shade pair is one model: for example dark orange and dark blue. All models have the same likelihood. Included extinction rate shapes are exponentially increasing, exponentially decreasing, and a modal extinction event. The speciation rate shapes are linear increases, an instantaneous up-shift, and an instantaneous down-shift. We also proposed rates that are randomly drawn from the horseshoe Markov random field (HSMRF) distribution.

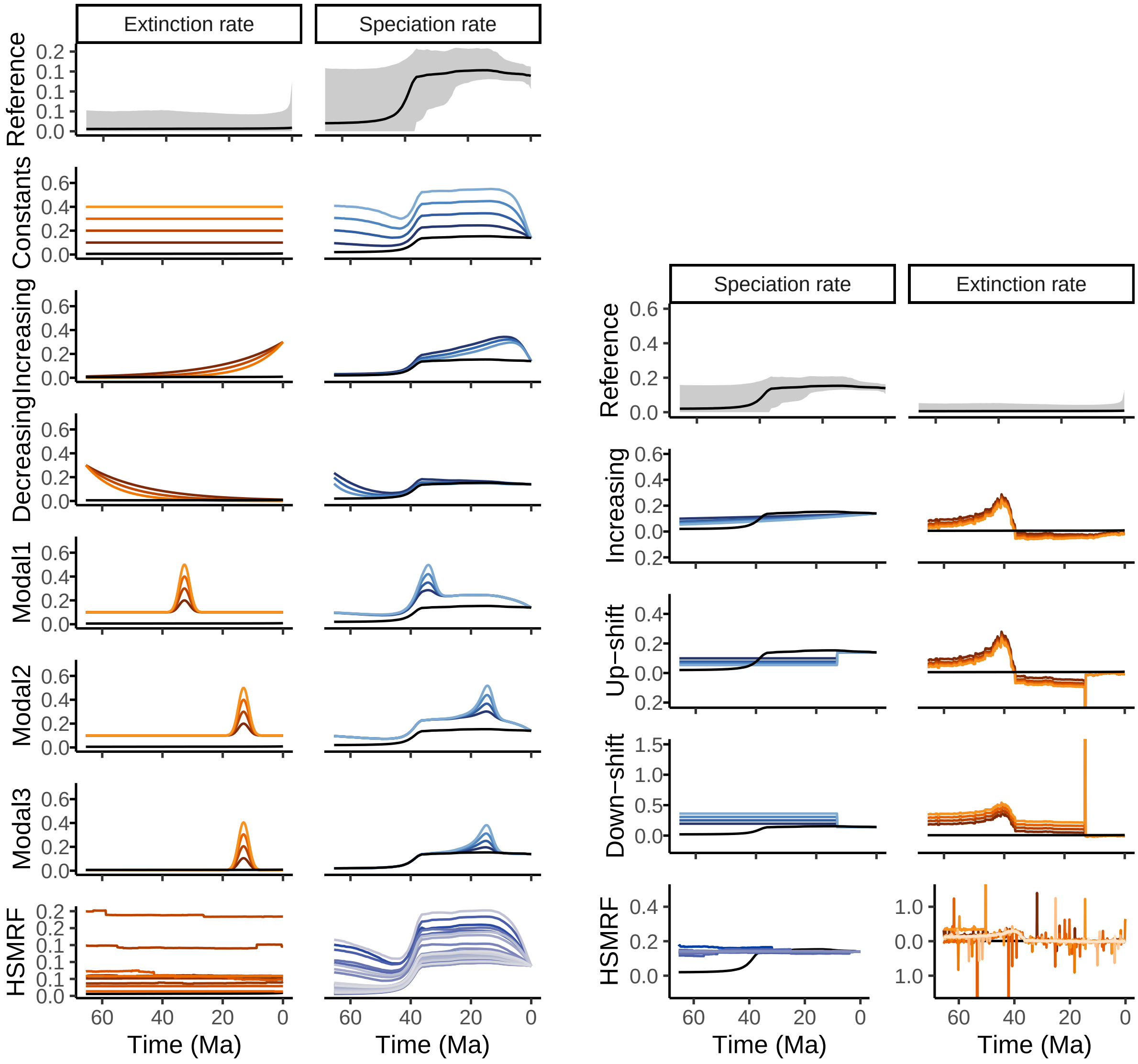

**Figure S5:** Congruent birth-death models for the tyrant flycatchers (Tyrannidae), based on a time-calibrated phylogenetic tree from (Condamine et al., 2019). Top row: the episodic birth-death model fitted in `RevBayes`, where the posterior median is in black, and the 95% credible interval in grey. The left panel exhibits models that are constructed by proposing alternative extinction rates. The right panel exhibits models that are constructed using alternative speciation rates. Each color-shade pair is one model: for example dark orange and dark blue. All models have the same likelihood. Included extinction rate shapes are exponentially increasing, exponentially decreasing, and a modal extinction event. The speciation rate shapes are linear increases, an instantaneous up-shift, and an instantaneous down-shift. We also proposed rates that are randomly drawn from the horseshoe Markov random field (HSMRF) distribution.

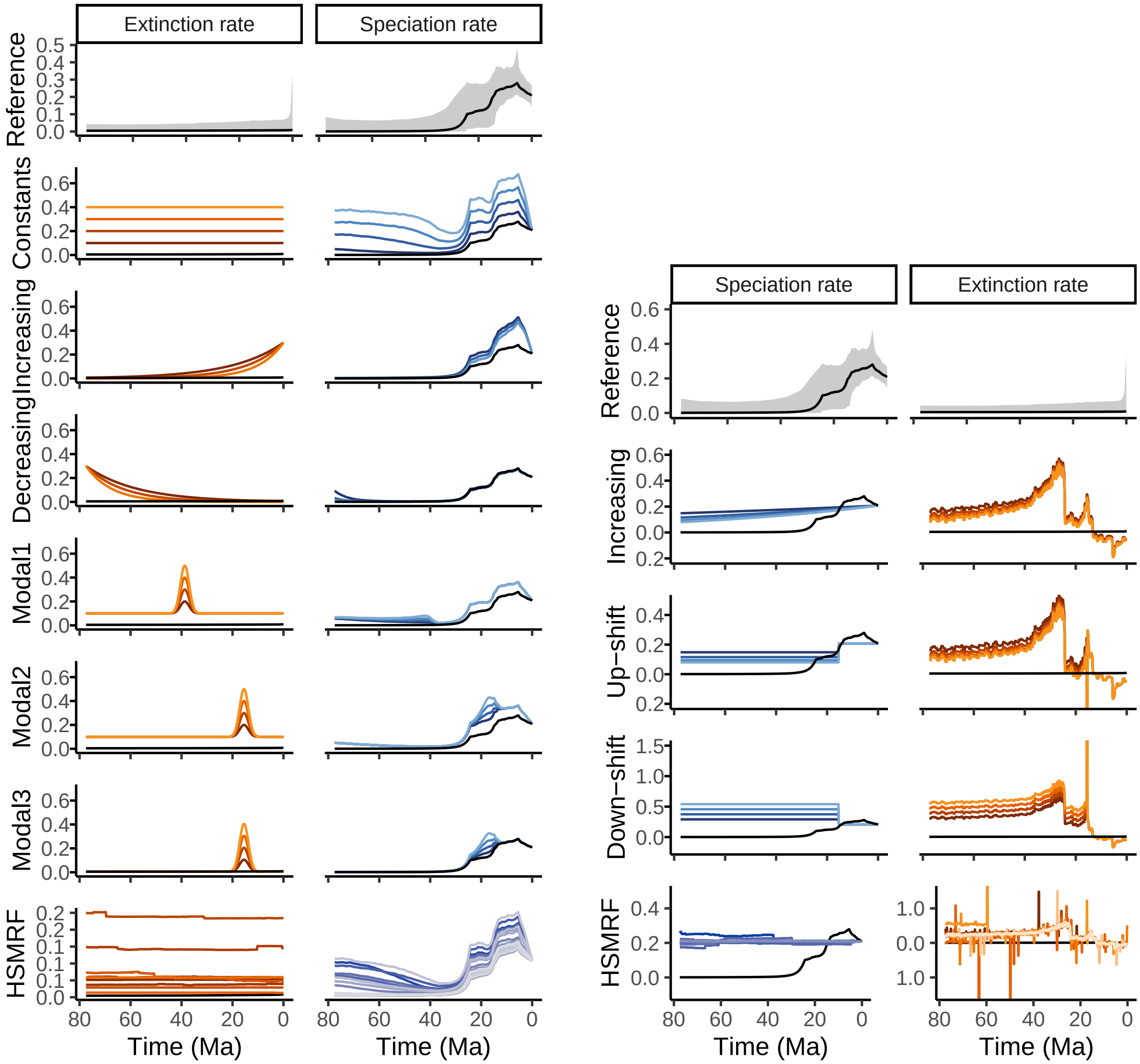

**Figure S6:** Congruent birth-death models for the tyrant woodpeckers (Picidae), based on a time-calibrated phylogenetic tree from Condamine et al. (2019). Top row: the episodic birth-death model fitted in `RevBayes`, where the posterior median is in black, and the 95% credible interval in grey. The left panel exhibits models that are constructed by proposing alternative extinction rates. The right panel exhibits models that are constructed using alternative speciation rates. Each color-shade pair is one model: for example dark orange and dark blue. All models have the same likelihood. Included extinction rate shapes are exponentially increasing, exponentially decreasing, and a modal extinction event. The speciation rate shapes are linear increases, an instantaneous up-shift, and an instantaneous down-shift. We also proposed rates that are randomly drawn from the horseshoe Markov random field (HSMRF) distribution.

### S3 Evaluating alternative trend thresholds

In the main text, and following Höhna et al. (2022), we proposed to assess the trends in the rate functions using the slope of the rate function. The slope is a continuous parameter, however, and can be a challenge to interpret. We chose to set a threshold of $\epsilon$ rate units per Ma, where if the slope was above the threshold, we considered the function to be significantly increasing (and analogously for decreasing rates). The choice of threshold is to some extent arbitrary. How much is a significant increase or decrease in the rate function? Choosing a threshold too small can result in picking up every small change (Fig. S7), and the trends will be difficult to interpret. Choosing a too large threshold (Fig. S9) will result in almost no changes being considered at all, and we are left with a blank, informationless figure. We recommend the researcher to test a few different thresholds that are suitable for their analysis, dataset, and relevant time scale.

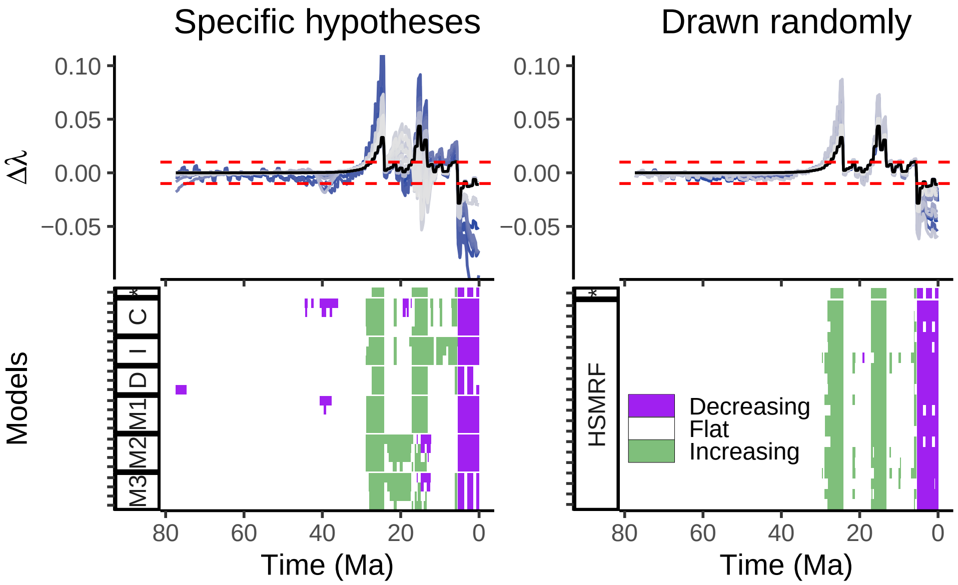

**Figure S7:** Congruent birth-death models for the tyrant woodpeckers (Picidae), based on a time-calibrated phylogenetic tree from Condamine et al. (2019), with a different threshold value $\epsilon$ to assess diversification rate trends. Top row: the slope of the speciation rate: $\Delta\lambda_i = (\lambda_i - \lambda_{i-1})/\Delta t$. The dashed red line indicates the threshold value of $\epsilon = 0.01$ rate units per Ma. Bottom row: a summary of whether each $\Delta\lambda$ is decreasing, increasing, or flat, as indicated by the threshold value. The extinction rate names are shortened, C: Constants, I: Increasing, D: Decreasing, M1-M3: Modals, HSMRF: horseshoe Markov random field, *: the reference model.

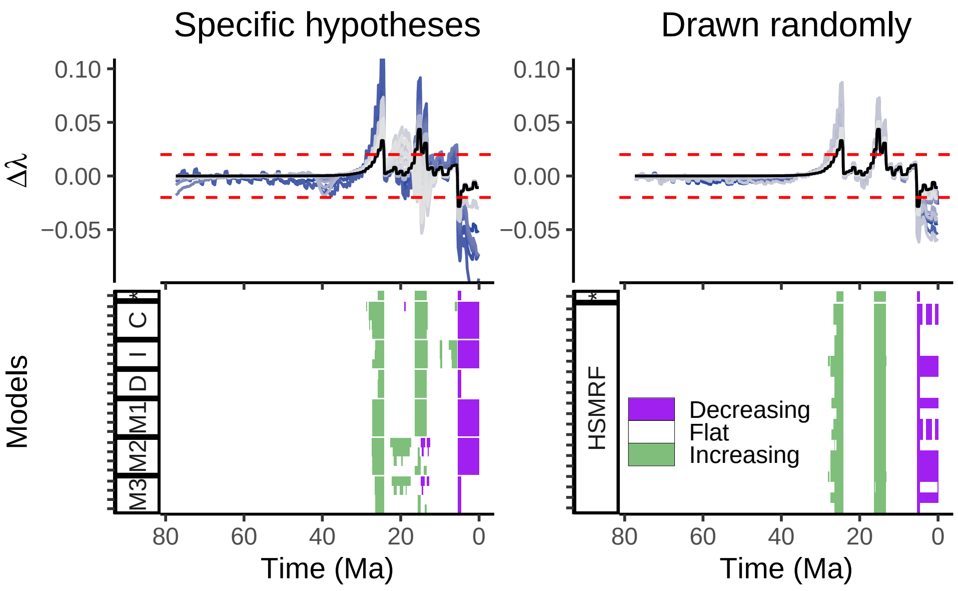

**Figure S8:** Equivalent to Fig. S7, however with a different threshold of $\epsilon = 0.02$ rate units per Ma. This threshold is the one we used for all other analyses and figures, unless otherwise specified.

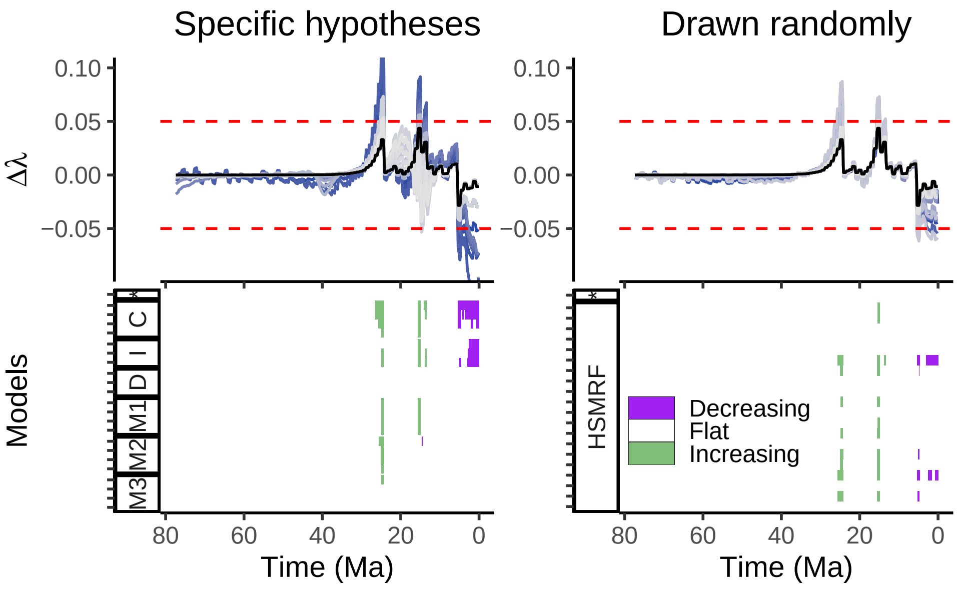

**Figure S9:** Equivalent to Fig. S7 and Fig. S8, however with a different threshold of $\epsilon = 0.05$ rate units per Ma. We did not use this threshold for any other analyses.

#### S4 Posterior samples

In this section we plot the first twenty samples from the posterior distribution (Fig. S10) that we used for the analyses in the main text (Fig. 3). This gives an indication of how variable the main temporal patterns in the rates are. For example, we can see that the samples from the woodpecker (Picidae) analysis largely display a strong increase in speciation rate, but the timing of when the first shift happened is uncertain, between 35 and 15 Ma (Fig. S10, bottom left panel). For this reason, we used a larger temporal window to calculate the directional trends from the posterior-distribution analyses, as opposed to the analyses using the posterior median. We computed the directional trend as $\Delta\lambda_i = (\lambda_i - \lambda_{i-k})/(t_i - t_{i-k}))$, where $k$ is the window size. In Fig. S11 we show how different choices of window sizes ($k \in 10, 25, 55, 80$) impact how the directional trends are portrayed.

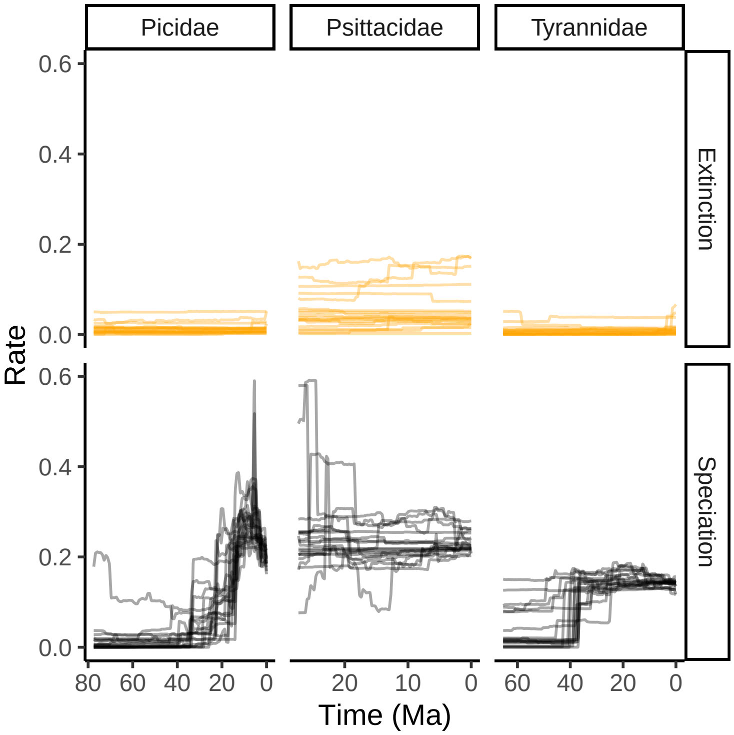

**Figure S10:** Speciation and extinction rate trajectories for the first twenty posterior samples used in assessing the trends in the congruence class when we take into consideration the uncertainty of the posterior estimates (main text Fig. 3).

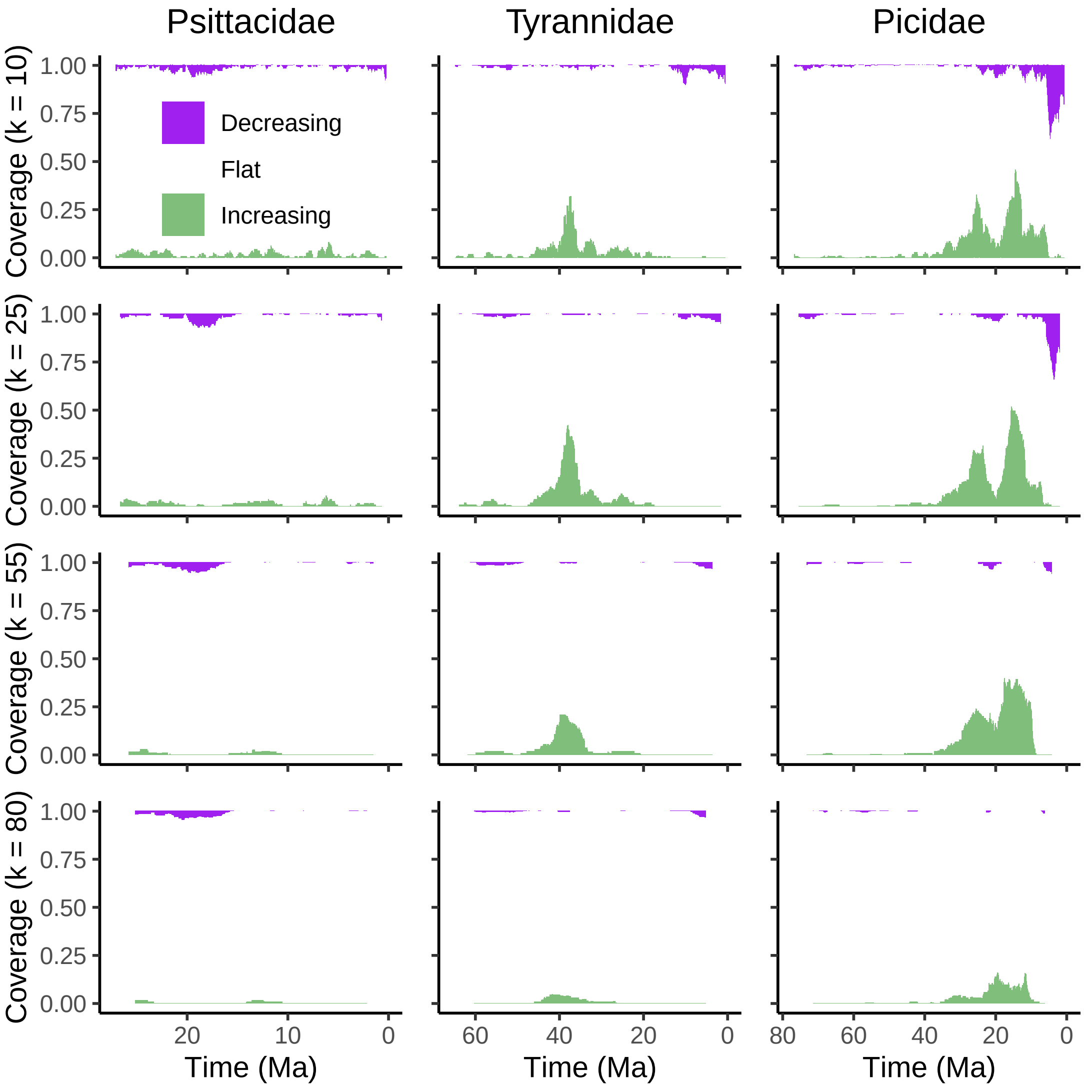

**Figure S11:** Directional trends in the speciation rate under 110 equidistant samples from the posterior distribution. Equal to main text Fig. 3, except with a variety of choices for the window size $k$, used when computing the slope in the speciation rate ($\Delta\lambda_i = (\lambda_i - \lambda_{i-k})/(t_i - t_{i-k})$)).

### S5 Convergence assessment

In this section we assess whether the Markov chain Monte Carlo (MCMC) simulation converged for the three datasets used in the main text: the parrots (Fig. S12), the tyrant flycatchers (Fig. S13), and the woodpeckers (Fig. S14). We plot the speciation and extinction rate estimates for each replicate run (Fig. S12 right, )(Fig. S13 right and (Fig. S14 right), and we perform the Kolmogorov-Smirnov (KS) test to assess whether the samples drawn from the posterior are reproducible across replicates. The KS-test is a measure of whether the samples could be considered to be drawn from the same underlying distribution, that is, the MCMC sampler has converged to the same posterior distribution (Fabreti and Höhna, 2022). For two equally distributed random variables, with 625 independent samples from each, we expect the KS-test to be false positive ($D > D_{\text{crit}} = 0.09$) in 1% of repeated comparisons (Fabreti and Höhna, 2022). Our computed KS scores are well below $D_{\text{crit}}$, which indicates that our samples have converged. The posterior median and credible intervals are similar to the across all four replicates for the woodpeckers, the tyrant flycatchers, and the parrots (Figs. S12 to S14).

For the analyses presented in the main text and otherwise in the supplementary materials, we pooled the four runs by concatenating the samples before computing the posterior medians and the credible intervals. This ensures more precise estimates in comparison to using each replicate individually. We conclude that our MCMC simulations converged towards the same parameter space and that our posterior median estimates are sufficiently precise for identifying the main time-varying patterns in the model.

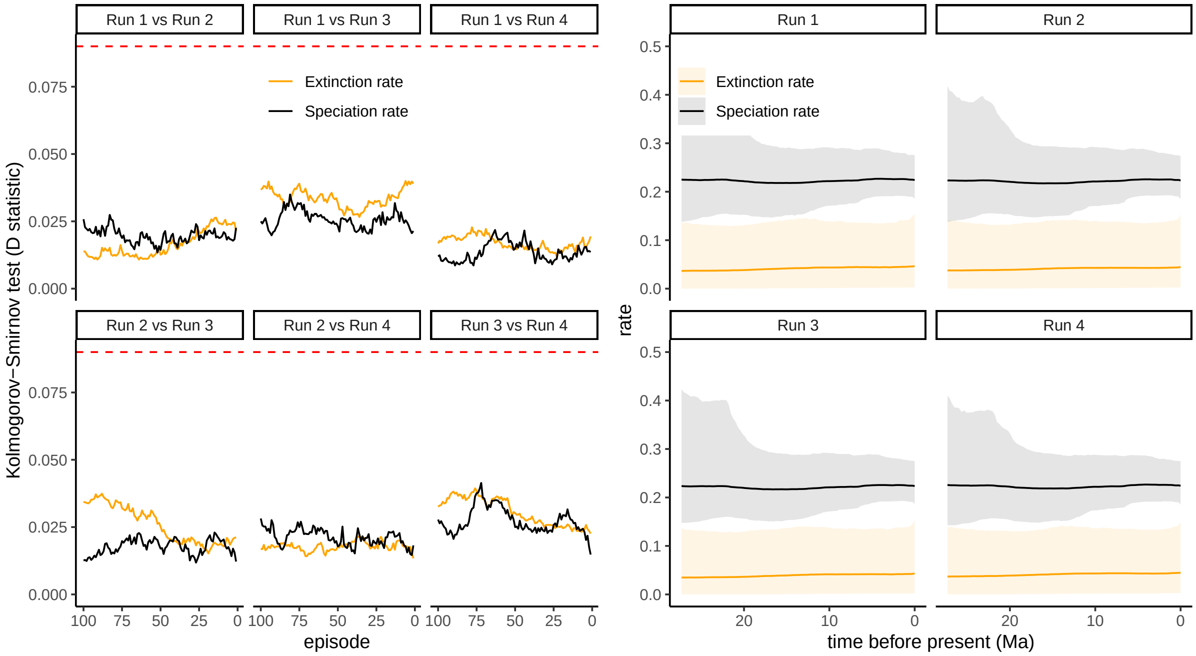

**Figure S12:** Convergence assessment for the New World/African true parrots, where we fitted an episodic birth-death model with a HSMRF (horseshoe Markov random field) distribution to model temporal autocorrelation in the rates. The analysis was performed in `RevBayes` with four replicates. We discarded the first 5000 iterations as burnin, ran the MCMC for another 50,000 iterations, and sampled every 10 iterations. The pairwise Kolmogorov-Smirnov test is an assessment of whether the two set of samples are drawn from the same distribution. Since we have $D < D_{\text{crit}} = 0.09$ (dashed red line), our replicates are similar, and the runs converged.

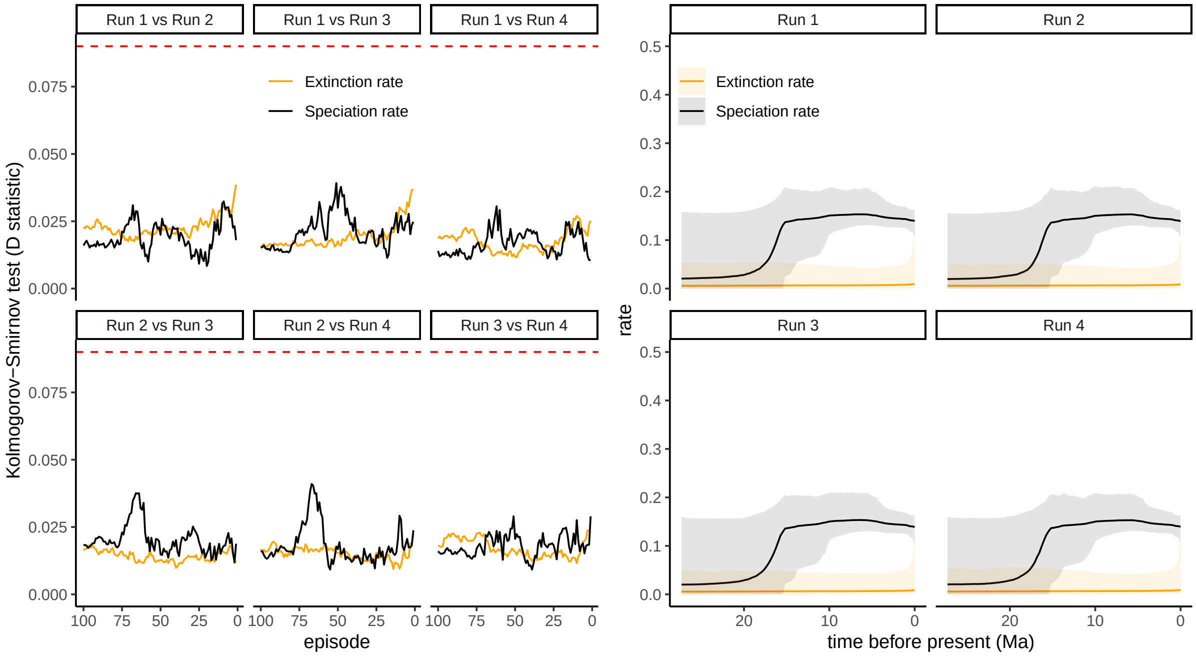

**Figure S13:** Convergence assessment for the tyrant flycatcher analysis, where we fitted an episodic birth-death model with a HSMRF (horseshoe Markov random field) distribution to model temporal autocorrelation in the rates. The analysis was performed in `RevBayes` with four replicates. We discarded the first 5000 iterations as burnin, ran the MCMC for another 50,000 iterations, and sampled every 10 iterations. The pairwise Kolmogorov-Smirnov test is an assessment of whether the two set of samples are drawn from the same distribution. Since we have $D < D_{\text{crit}} = 0.09$ (dashed red line), our replicates are similar, and the runs converged.

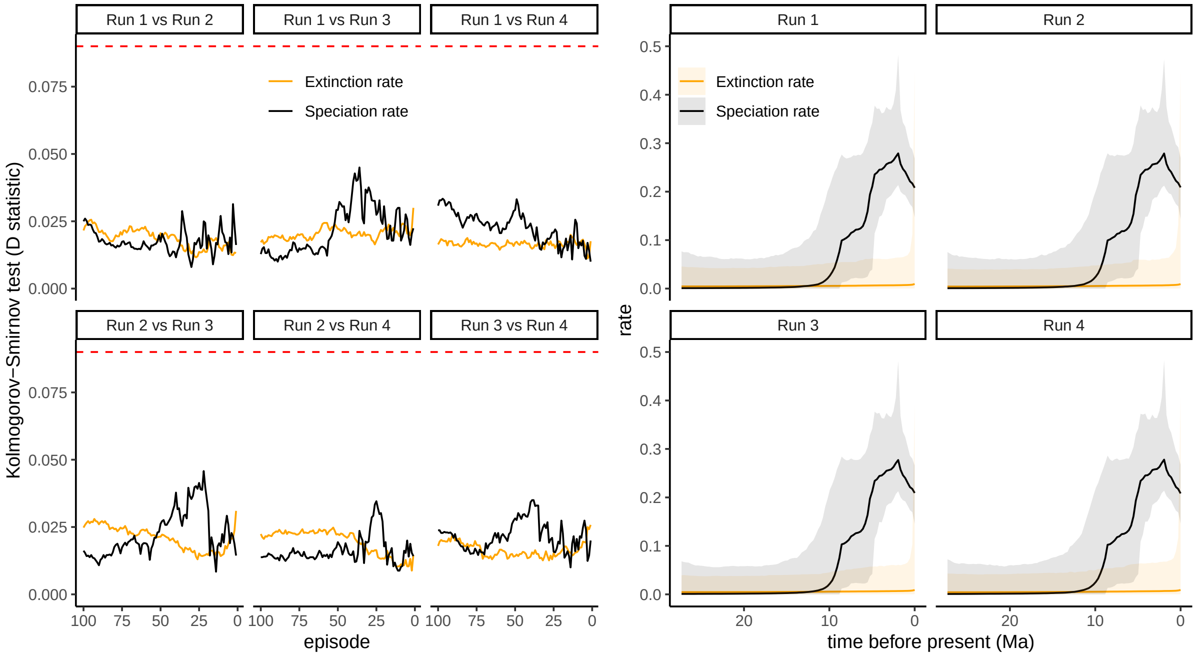

**Figure S14:** Convergence assessment for the woodpecker analysis, where we fitted an episodic birth-death model with a HSMRF (horseshoe Markov random field) distribution to model temporal autocorrelation in the rates. The analysis was performed in `RevBayes` with four replicates. We discarded the first 5000 iterations as burnin, ran the MCMC for another 50,000 iterations, and sampled every 10 iterations. The pairwise Kolmogorov-Smirnov test is an assessment of whether the two set of samples are drawn from the same distribution. Since we have $D < D_{\text{crit}} = 0.09$ (dashed red line), our replicates are similar, and the runs converged.

### S6 Hypothetical models: alternative extinction rate

In the main text (Fig. 2) we showed three hypothetical diversification histories, and we proposed eight alternative categories of rate shapes that we used to construct congruent models. In this section we expand the three classes to eight sets of congruent models (Fig. S15). In all cases we used a constant extinction rate of $\mu = 0.28$. These congruent models exhibit a variety of inferred speciation rate trajectories, which we depict in this section (Figs. S16 to S23). For all datasets that show a rapid change in the speciation rate in the reference model, then all alternative models also show the same rapid change in speciation rates as shown by our assessment of directional trends. Conversely, if the reference speciation rate does not show a directional trend (i.e., the rate is approximately constant), then either the alternative speciation rates also showed no directional trend or the trends among alternative models were in disagreement.

We observed the strongest disagreement of trends for the speciation rate intervals near the present, especially for the constant reference model. The conflicting trends correspond directly to the scale of the proposed extinction rate relative to the reference speciation rate ($\lambda = 0.35$, panel C). The proposed extinction rates that are larger than the reference extinction rate $\mu = 0.28$ at the near-present (rates L+, S+, E+, and some constant rate functions) infer models with decreasing speciation rates near the present (Fig. S15, panel C and Fig. S16). Conversely, proposed extinction rates that are at present smaller than $\mu = 0.28$ infer models with increasing speciation rates near the present (Fig. S15, panel C and Fig. S16). Similar patterns can be seen at the near-present for all of the other congruence classes (Figs. S17 to S23). As we observed in our empirical datasets, if the reference speciation rate model was constant, then we observe no fully supported trends in the congruence class (Fig. S15, panel C). We see that models with moderate speciation rate changes do not show shared patterns across the congruence class (panels L+, L–). Reference models with rapidly changing rates show unambiguous agreement for their correspond trend at the shift time (panels S+, S–). The robustness of the trends regarding the congruence class weakens when the rate change is less steep (panels M, E+, E–).

The overall results of the hypothetical cases confirm the conclusions from the empirical datasets: changes that happen slowly over time are not robust to the congruence class. However, changes that happen abruptly in a small time window are indeed robust.

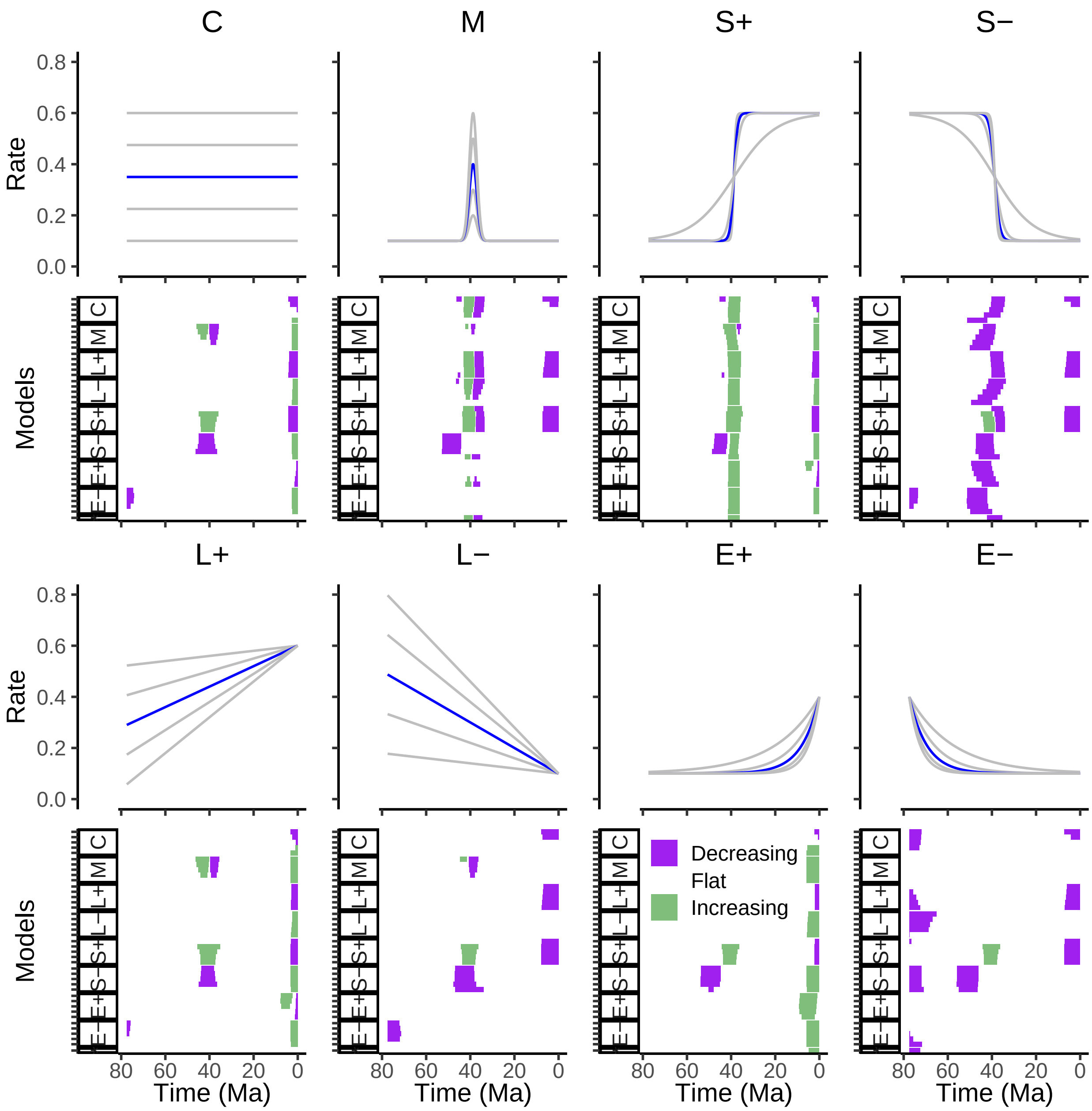

**Figure S15:** A collection of hypothetical scenarios. Each panel-pair represents a separate congruence class. The reference model for the congruence class is the speciation rate depicted in blue, and in all cases a constant extinction rate of $\mu = 0.28$. For each congruence class, we proposed various alternative extinction rates: constants (C), modals (M), sigmoidal increase (S+), sigmoidal decrease (S–), linear increase (L+), linear decrease (L–), exponential increase (E+) and exponential decrease (E–). Each congruence class results in a set of models with inferred speciation rate curves (Figs. S16 to S23). We computed the significant directional trends for the inferred speciation rates (with a threshold of $\epsilon = 0.02$ rate units per Ma) for each congruence class. We selected a time scale similar to the woodpecker phylogeny for interpretability.

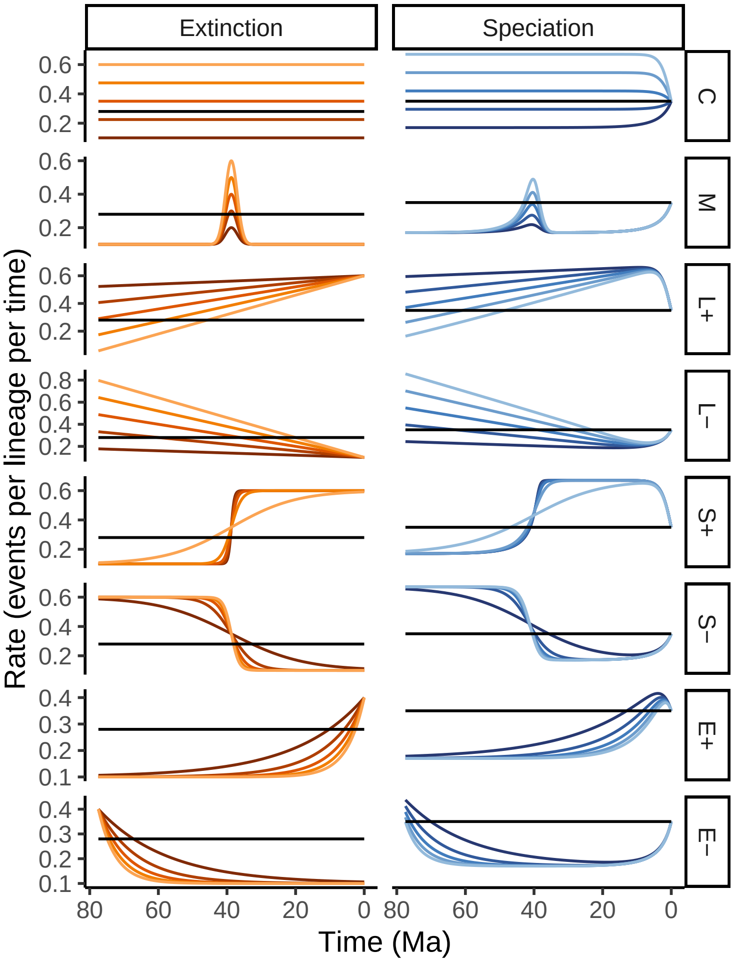

**Figure S16:** Hypothetical rate scenarios for the constant birth-death model in Fig. S15. The reference model is depicted in black in all panels: a constant extinction rate ($\mu = 0.28$), and a constant speciation rate function corresponding to ( Fig. S15, C, red line). Five alternative extinction rates are proposed in each row, including constants (C), modals (M), linear increase (L+), linear decrease (L–), sigmoidal increase (S+), sigmoidal decrease (S–), exponential increase (E+), and exponential decrease (E–). The right column depicts the congruent speciation rates.

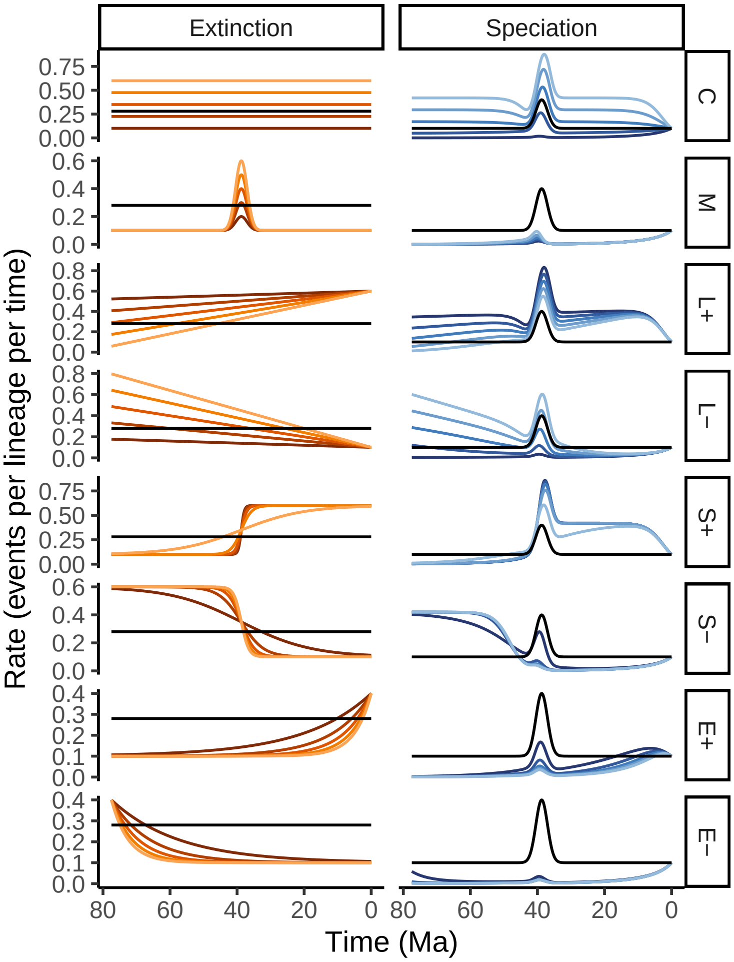

**Figure S17:** Hypothetical rate scenarios for the modal birth-death model in Fig. S15. The reference model is depicted in black in all panels: a constant extinction rate ($\mu = 0.28$), and a modal speciation rate function corresponding to (Fig. S15, M, red line). Five alternative extinction rates are proposed in each row, including constants (C), modals (M), linear increase (L+), linear decrease (L–), sigmoidal increase (S+), sigmoidal decrease (S–), exponential increase (E+), and exponential decrease (E–). The right column depicts the congruent speciation rates.

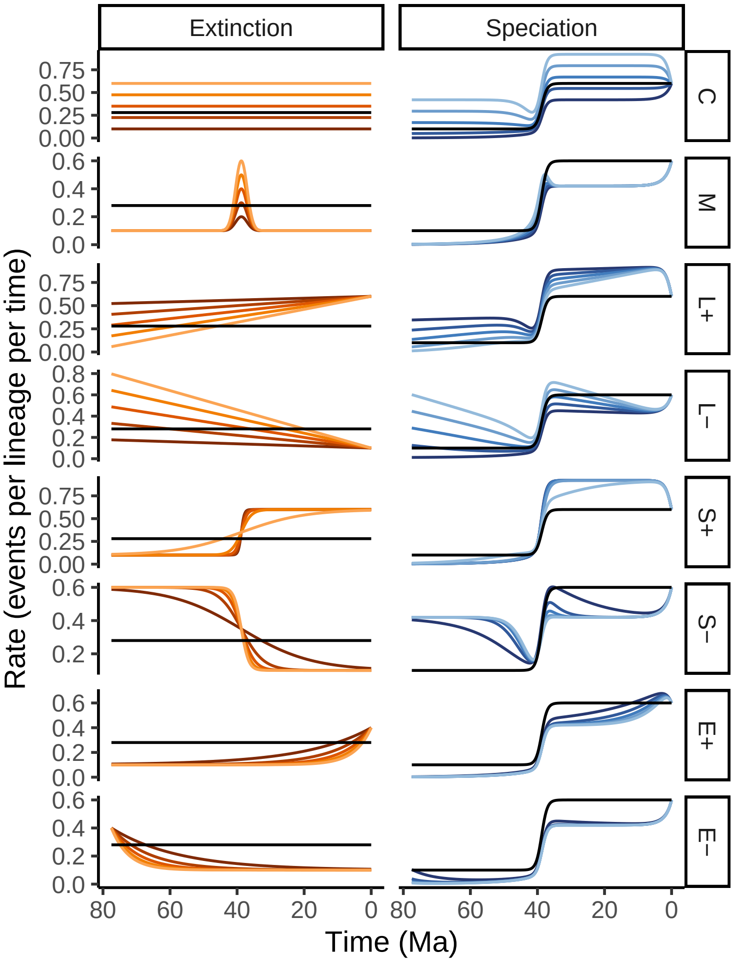

**Figure S18:** Hypothetical rate scenarios for the sigmoidally increasing birth-death model in Fig. S15. The reference model is depicted in black in all panels: a constant extinction rate ($\mu = 0.28$), and a sigmoidally increasing speciation rate function corresponding to ( Fig. S15, S+, red line). Five alternative extinction rates are proposed in each row, including constants (C), modals (M), linear increase (L+), linear decrease (L–), sigmoidal increase (S+), sigmoidal decrease (S–), exponential increase (E+), and exponential decrease (E–). The right column depicts the congruent speciation rates.

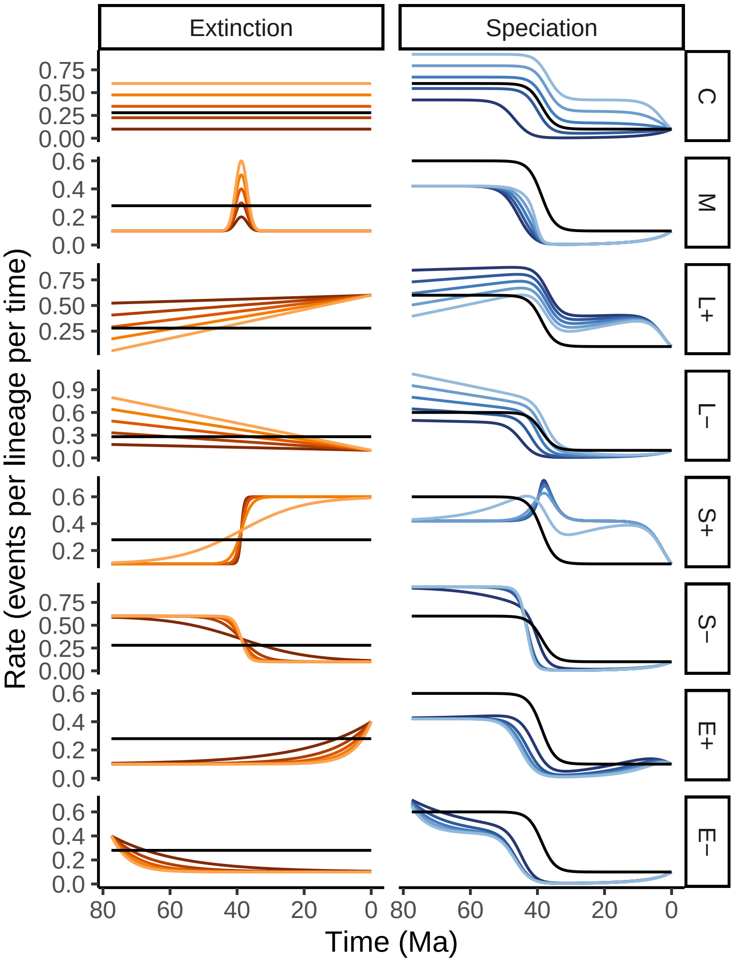

**Figure S19:** Hypothetical rate scenarios for the sigmoidally decreasing birth-death model in Fig. S15. The reference model is depicted in black in all panels: a constant extinction rate ($\mu = 0.28$), and a sigmoidally decreasing speciation rate function corresponding to ( Fig. S15, S–, red line). Five alternative extinction rates are proposed in each row, including constants (C), modals (M), linear increase (L+), linear decrease (L–), sigmoidal increase (S+), sigmoidal decrease (S–), exponential increase (E+), and exponential decrease (E–). The right column depicts the congruent speciation rates.

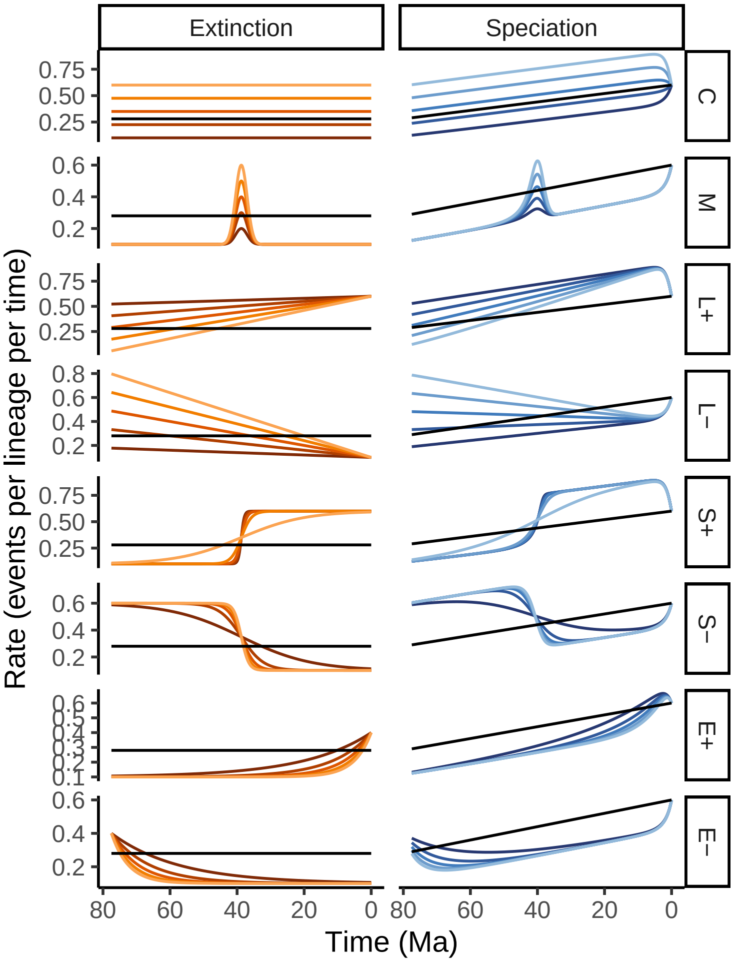

**Figure S20:** Hypothetical rate scenarios for the linearly increasing birth-death model in Fig. S15. The reference model is depicted in black in all panels: a constant extinction rate ($\mu = 0.28$), and a linearly increasing speciation rate function corresponding to ( Fig. S15, L+, red line). Five alternative extinction rates are proposed in each row, including constants (C), modals (M), linear increase (L+), linear decrease (L–), sigmoidal increase (S+), sigmoidal decrease (S–), exponential increase (E+), and exponential decrease (E–). The right column depicts the congruent speciation rates.

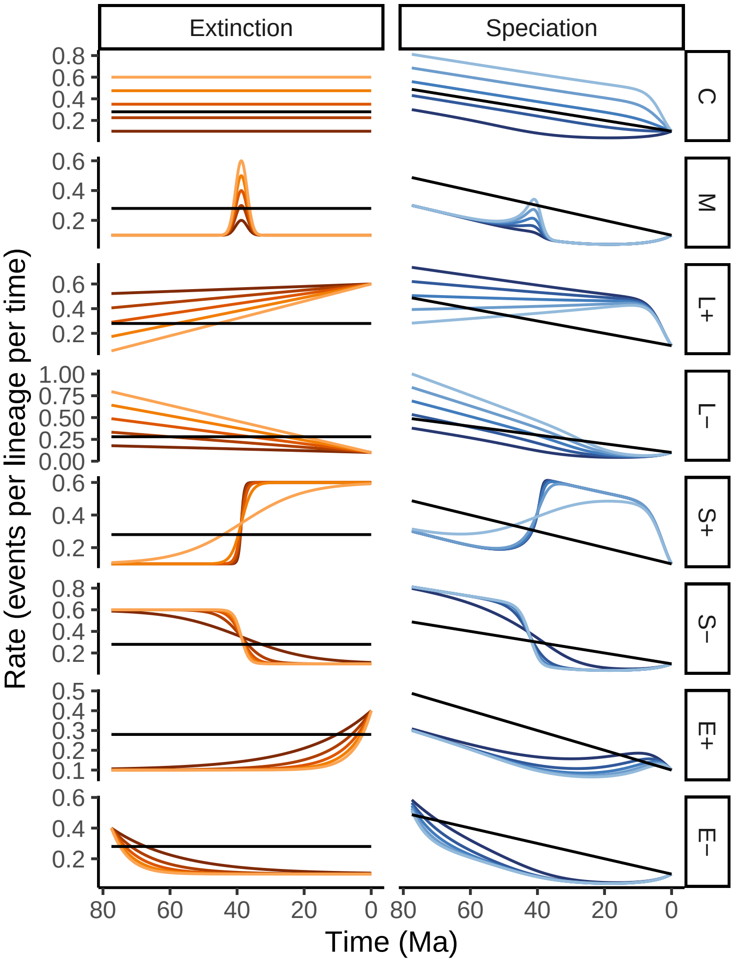

**Figure S21:** Hypothetical rate scenarios for the linearly decreasing birth-death model in Fig. S15. The reference model is depicted in black in all panels: a constant extinction rate ($\mu = 0.28$), and a linearly decreasing speciation rate function corresponding to ( Fig. S15, L–, red line). Five alternative extinction rates are proposed in each row, including constants (C), modals (M), linear increase (L+), linear decrease (L–), sigmoidal increase (S+), sigmoidal decrease (S–), exponential increase (E+), and exponential decrease (E–). The right column depicts the congruent speciation rates.

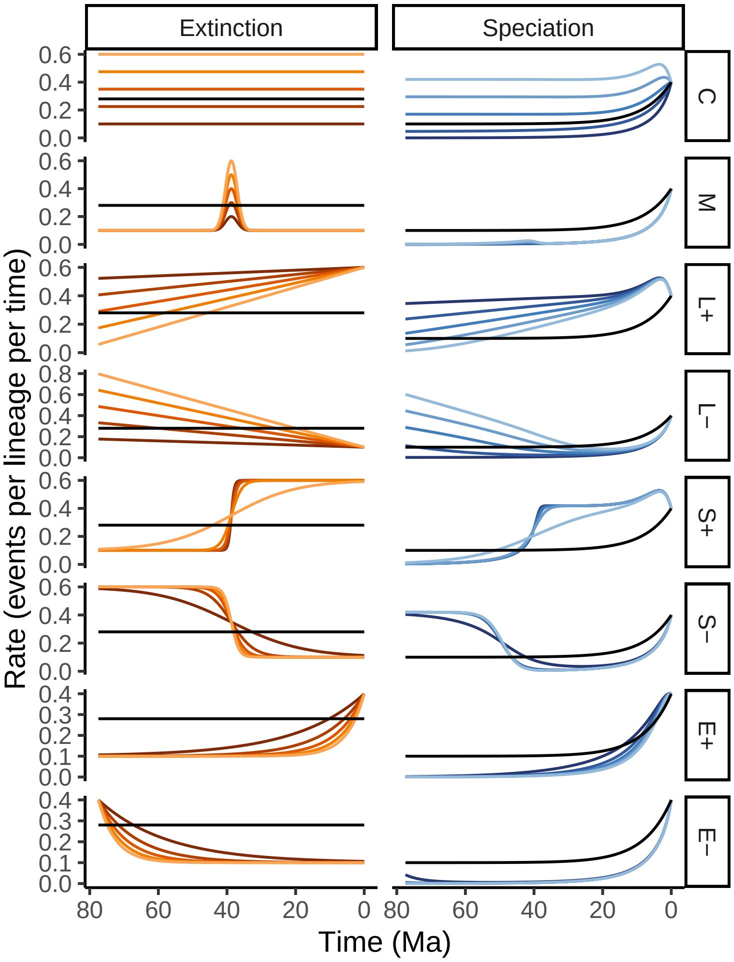

**Figure S22:** Hypothetical rate scenarios for the exponentially increasing birth-death model in Fig. S15. The reference model is depicted in black in all panels: a constant extinction rate ($\mu = 0.28$), and an exponentially increasing speciation rate function corresponding to ( Fig. S15, E+, red line). Five alternative extinction rates are proposed in each row, including constants (C), modals (M), linear increase (L+), linear decrease (L–), sigmoidal increase (S+), sigmoidal decrease (S–), exponential increase (E+), and exponential decrease (E–). The right column depicts the congruent speciation rates.

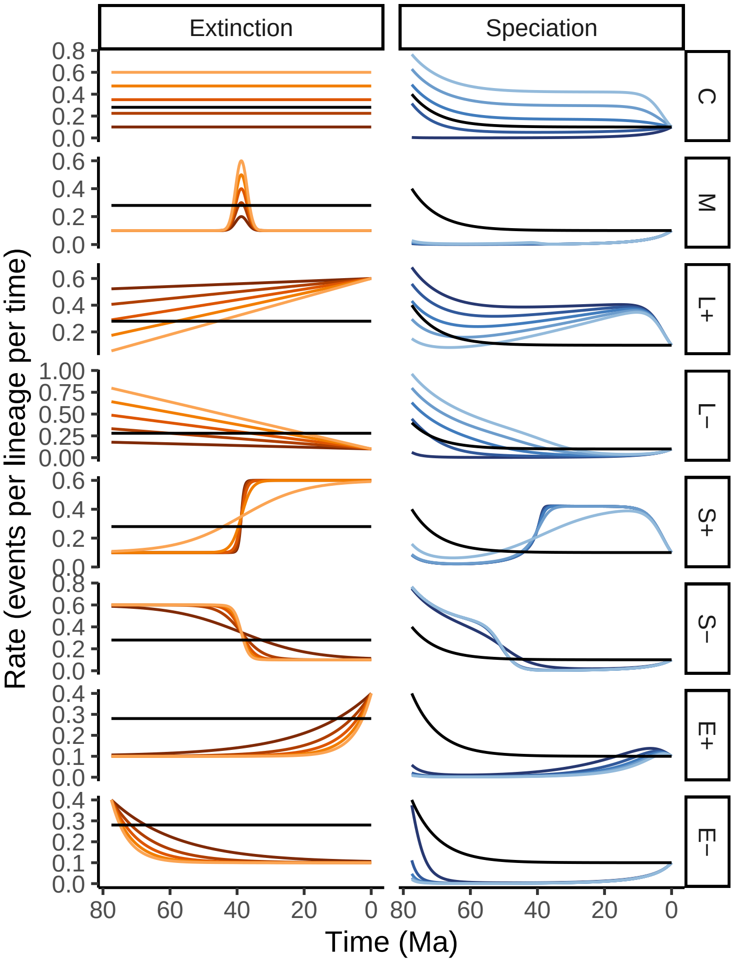

**Figure S23:** Hypothetical rate scenarios for the exponentially decreasing birth-death model in Fig. S15. The reference model is depicted in black in all panels: a constant extinction rate ($\mu = 0.28$), and an exponentially decreasing speciation rate function corresponding to ( Fig. S15, E-, red line). Five alternative extinction rates are proposed in each row, including constants (C), modals (M), linear increase (L+), linear decrease (L–), sigmoidal increase (S+), sigmoidal decrease (S–), exponential increase (E+), and exponential decrease (E–). The right column depicts the congruent speciation rates.

### S7 Hypothetical models: alternative speciation rate

In the previous section, we explored hypothetical models where we proposed alternative extinction rate functions. In this section, we set up hypothetical models where we instead propose alternative speciation rate functions. As discussed in the main text, there is an additional constraint to be considered when proposing speciation rate functions. Namely, the speciation rate at the present ($\lambda_0$) must be equal for all models in the congruence class. Since we could not vary the constant-rate model with different intercepts according to this constraint, we omitted this group of rate functions. In all seven congruence classes, we constructed a reference with constant speciation rate ($\lambda = 0.28$), and for the extinction rate the red trajectories seen in (Fig. S24). Next, we proposed a variety of speciation rate trajectories, including a modal event (Fig. S25), sigmoidally increasing (Fig. S26), sigmoidally decreasing (Fig. S27), linearly increasing (Fig. S28), linearly decreasing (Fig. S29), exponentially increasing (Fig. S30), and exponentially decreasing (Fig. S31).

In comparison to our empirical examples, we were able to propose more valid alternative speciation rate functions (that did not result in negative extinction rate functions). This may be due to our reference extinction rates (Fig. S24, red lines) being larger than those estimated from our empirical datasets, and thus there is more room for variation in the extinction rate before it goes negative. Note that we used a constant reference speciation rate, and because all models have the same $\lambda_0$, the linear and exponentially increasing alternative speciation rate functions (L+ and E+) as well as the up-shift (sigmoidal, S+) produce necessarily lower alternative speciation rate functions.

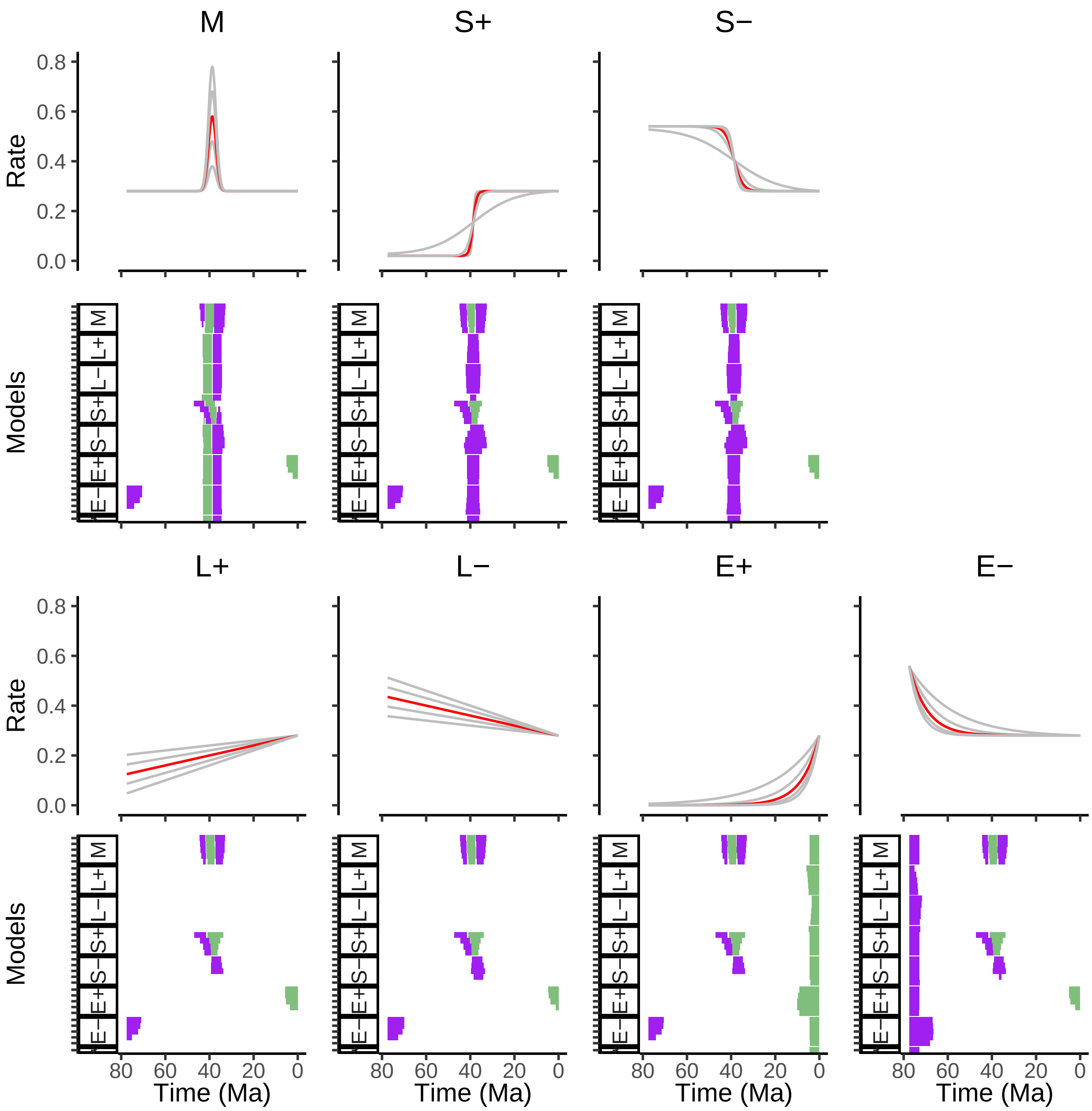

**Figure S24:** A collection of hypothetical models. Each panel-pair represents a separate congruence class. The reference model for the congruence class is the extinction rate depicted in red, and in all cases a constant speciation rate of $\lambda = 0.28$. For each congruence class, we proposed various alternative speciation rates: modals (M), linear increase (L+), linear decrease (L–), sigmoidal increase (S+), sigmoidal decrease (S–), exponential increase (E+) and exponential decrease (E–). Each congruence class results in a set of models with inferred speciation rate curves (See Figs. S25 to S31). We computed the significant directional trends for the inferred extinction rates (with a threshold of $\epsilon = 0.02$ rate units per Ma) for each congruence class. We selected a time scale similar to the woodpecker phylogeny for interpretability.

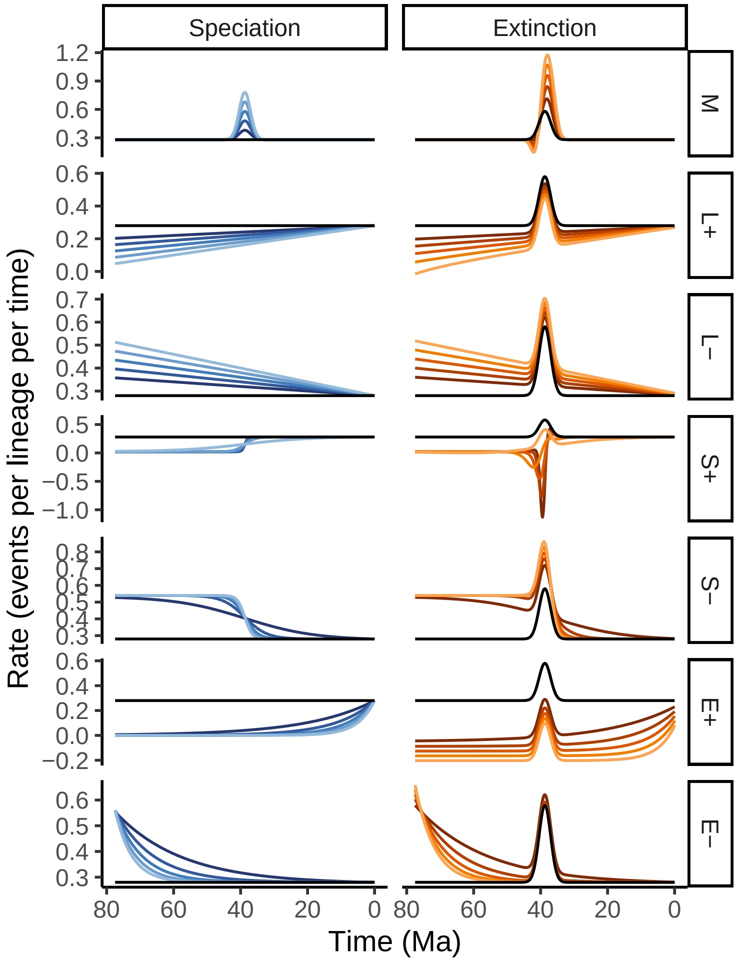

**Figure S25:** Hypothetical rate scenarios for the modal birth-death model in Fig. S24. The reference model is depicted in black in all panels: a constant speciation rate ($\lambda = 0.28$), and a modal extinction rate function corresponding to (Fig. S24, M, red line). Five alternative speciation rates are proposed in each row, including modals (M), linear increase (L+), linear decrease (L–), sigmoidal increase (S+), sigmoidal decrease (S–), exponential increase (E+), and exponential decrease (E–). The right column depicts the congruent extinction rates.

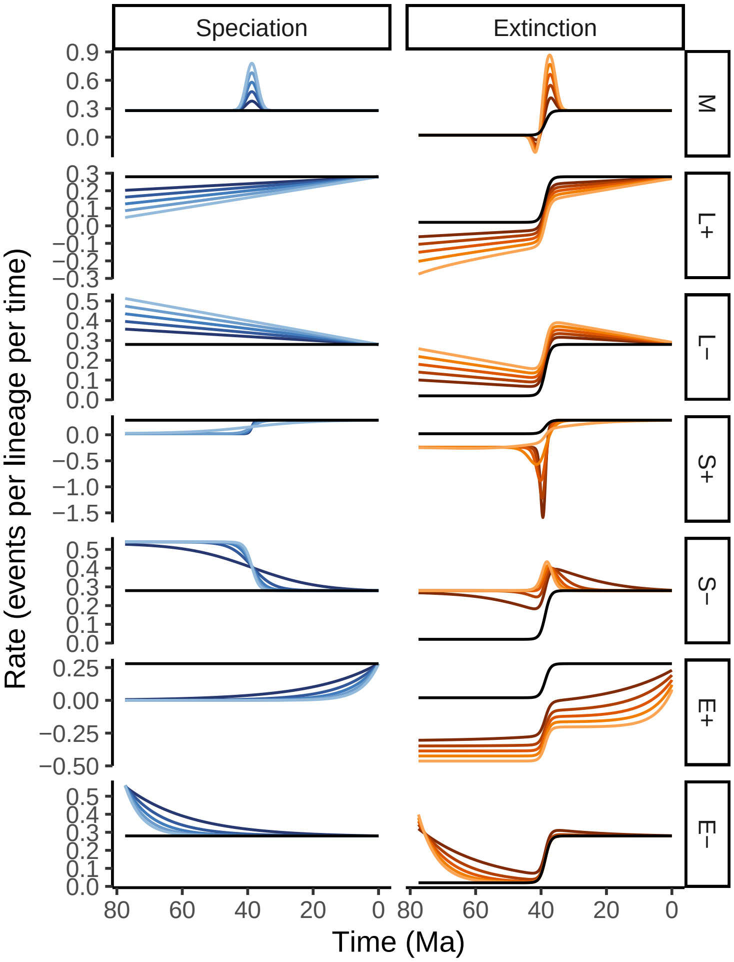

**Figure S26:** Hypothetical rate scenarios for the sigmoidally increasing birth-death model in Fig. S24. The reference model is depicted in black in all panels: a constant speciation rate ($\lambda = 0.28$), and a sigmoidally increasing extinction rate function corresponding to (Fig. S24, S+, red line). Five alternative speciation rates are proposed in each row, including modals (M), linear increase (L+), linear decrease (L–), sigmoidal increase (S+), sigmoidal decrease (S–), exponential increase (E+), and exponential decrease (E–). The right column depicts the congruent extinction rates.

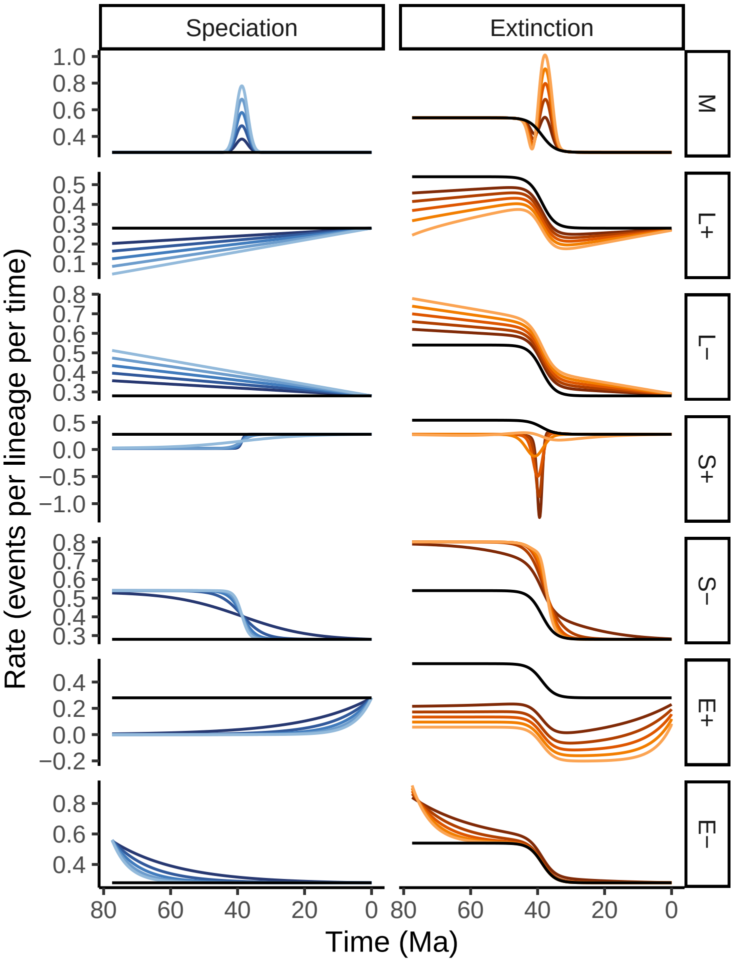

**Figure S27:** Hypothetical rate scenarios for the sigmoidally decreasing birth-death model in Fig. S24. The reference model is depicted in black in all panels: a constant speciation rate ($\lambda = 0.28$), and a sigmoidally decreasing extinction rate function corresponding to (Fig. S24, S–, red line). Five alternative speciation rates are proposed in each row, including modals (M), linear increase (L+), linear decrease (L–), sigmoidal increase (S+), sigmoidal decrease (S–), exponential increase (E+), and exponential decrease (E–). The right column depicts the congruent extinction rates.

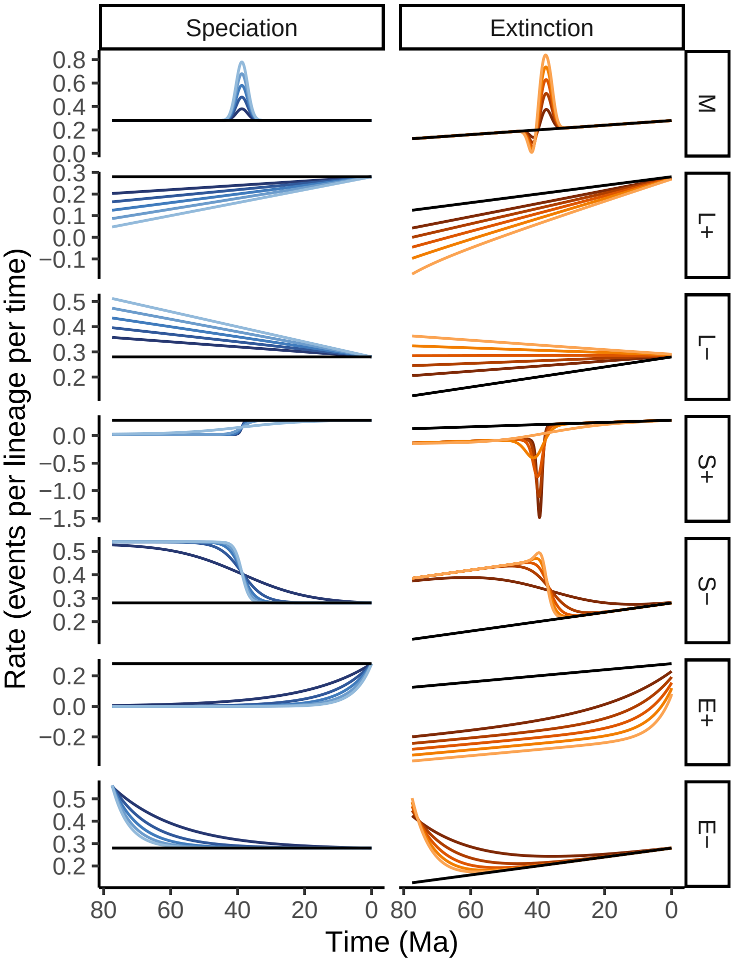

**Figure S28:** Hypothetical rate scenarios for the linearly increasing birth-death model in Fig. S24. The reference model is depicted in black in all panels: a constant speciation rate ($\lambda$ = 0.28), and a linearly increasing extinction rate function corresponding to (Fig. S24, L+, red line). Five alternative speciation rates are proposed in each row, including modals (M), linear increase (L+), linear decrease (L–), sigmoidal increase (S+), sigmoidal decrease (S–), exponential increase (E+), and exponential decrease (E–). The right column depicts the congruent extinction rates.

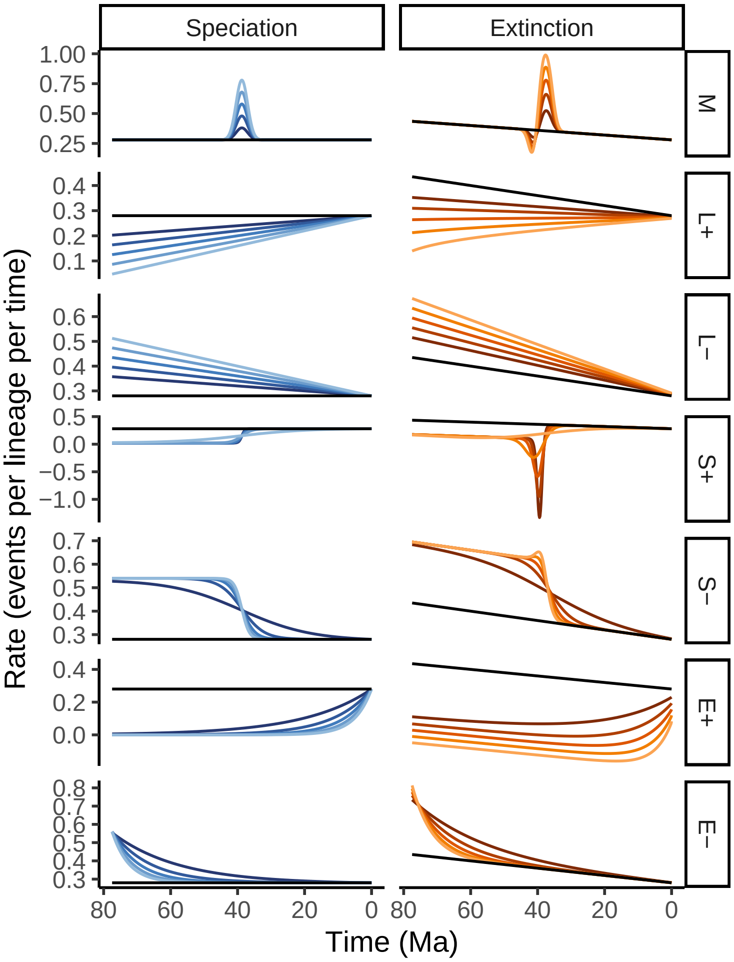

**Figure S29:** Hypothetical rate scenarios for the linearly decreasing birth-death model in Fig. S24. The reference model is depicted in black in all panels: a constant speciation rate ($\lambda = 0.28$), and a linearly decreasing extinction rate function corresponding to (Fig. S24, L–, red line). Five alternative speciation rates are proposed in each row, including modals (M), linear increase (L+), linear decrease (L–), sigmoidal increase (S+), sigmoidal decrease (S–), exponential increase (E+), and exponential decrease (E–). The right column depicts the congruent extinction rates.

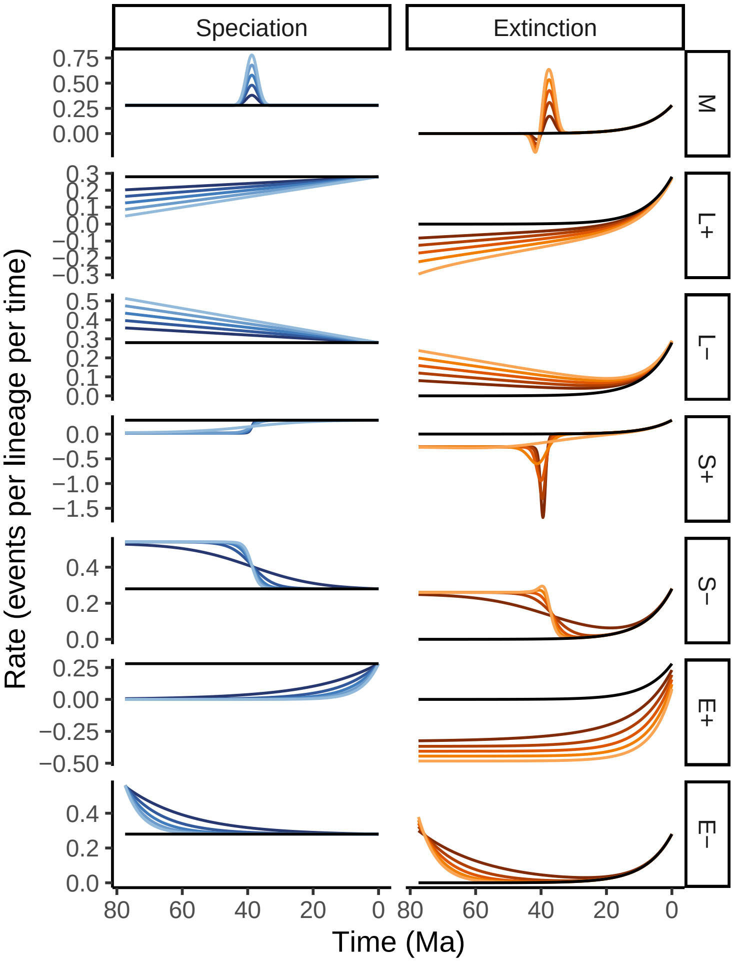

**Figure S30:** Hypothetical rate scenarios for the exponentially increasing birth-death model in Fig. S24. The reference model is depicted in black in all panels: a constant speciation rate ($\lambda = 0.28$), and an exponentially increasing extinction rate function corresponding to (Fig. S24, E+, red line). Five alternative speciation rates are proposed in each row, including modals (M), linear increase (L+), linear decrease (L–), sigmoidal increase (S+), sigmoidal decrease (S–), exponential increase (E+), and exponential decrease (E–). The right column depicts the congruent extinction rates.

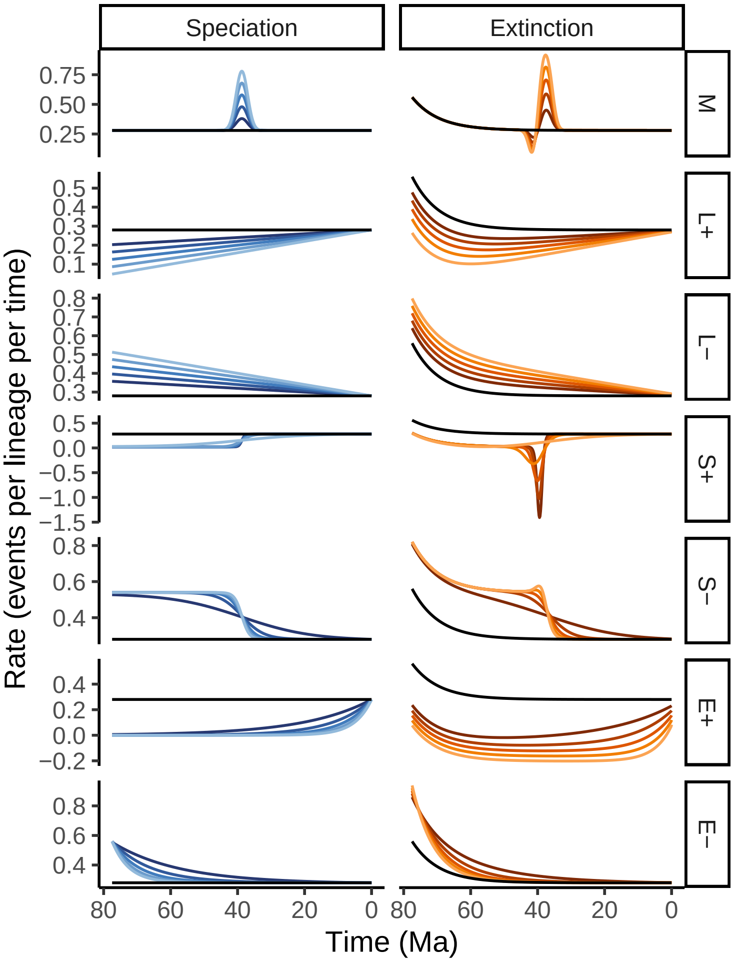

**Figure S31:** Hypothetical rate scenarios for the exponentially decreasing birth-death model in Fig. S24. The reference model is depicted in black in all panels: a constant speciation rate ($\lambda = 0.28$), and an exponentially decreasing extinction rate function corresponding to (Fig. S24, E–, red line). Five alternative speciation rates are proposed in each row, including modals (M), linear increase (L+), linear decrease (L–), sigmoidal increase (S+), sigmoidal decrease (S–), exponential increase (E+), and exponential decrease (E–). The right column depicts the congruent extinction rates.
